# Independent regulation of mtDNA quantity and quality resets the mitochondrial genome in *C. elegans* primordial germ cells

**DOI:** 10.1101/2022.05.06.490954

**Authors:** Aaron Z.A. Schwartz, Nikita Tsyba, Yusuff Abdu, Maulik R. Patel, Jeremy Nance

## Abstract

Mitochondria contain an independent genome, called mtDNA, which contains essential metabolic genes. Although mtDNA mutations occur at high frequency, they are inherited infrequently, indicating that germline mechanisms limit their accumulation. To determine how germline mtDNA is regulated, we examined the control of mtDNA quantity and quality in *C. elegans* primordial germ cells (PGCs). We show that PGCs generate a bottleneck in mtDNA number by segregating mitochondria into lobe-like protrusions that are cannibalized by adjacent cells, reducing mtDNA content two-fold. As PGCs exit quiescence and divide, mtDNAs replicate to maintain a set point of ∼200 mtDNAs per germline stem cell. Whereas PGC lobe cannibalism eliminates mtDNAs stochastically, we show that the kinase PINK1, operating independently of Parkin and autophagy, preferentially reduces the fraction of mutant mtDNAs. Thus, PGCs employ parallel mechanisms to control both the quantity and quality of the founding population of germline mtDNAs.

## Introduction

Mitochondria contain multiple copies of a small genome called mitochondrial DNA (mtDNA), which contains several genes essential for oxidative phosphorylation [1]. Compared to nuclear DNA, mtDNA has a high mutation rate and is repaired inefficiently [1]. mtDNAs containing deleterious mutations are found together with complementing wild-type mtDNAs in a state called heteroplasmy. Deleterious mtDNA mutations can lead to mitochondrial disease if present at sufficiently high heteroplasmy – a condition that is estimated to affect ∼1 in 5000 individuals and has no known cure [2].

mtDNA replicates independently from nuclear DNA and has a distinct mode of inheritance. During cell division in most cell types, each daughter cell inherits a stochastic subset of mitochondria and their mtDNAs. However, embryos inherit their mtDNAs exclusively from the pool present within the oocyte [3]. The strict maternal inheritance and high mutation rate of mtDNA raise a potential problem: mtDNA mutations could accumulate over generations, leading to mutational meltdown [4]. However, relatively few deleterious mutations are transmitted over generations [5], indicating that mtDNA mutations are selected against within the germ line.

Two mechanisms have been proposed to regulate germline mtDNA inheritance. In one mechanism – the mitochondrial bottleneck – mtDNAs are reduced in number within the germline lineage to create a small founding population, which is passed on to the next generation. Bottlenecks are theorized to allow for the stochastic enrichment or depletion of variant mtDNAs [3, 6, 7]. In vertebrates, a bottleneck occurs in embryonic primordial germ cells (PGCs) due to the dilution of maternally provided mtDNAs by reductive embryonic cell divisions, or via the replication of a subset of mtDNA genomes in PGCs [8–13]. It is not known whether germline mtDNA bottlenecks could form through other means.

Alternatively, mitochondria containing high levels of mutant mtDNAs can be eliminated directly from germ cells – a process called purifying selection [3]. The mechanistic basis for germline purifying selection has been studied most intensively in the *Drosophila* ovary, where mtDNA mutations are eliminated both by autophagy and selective mtDNA replication [14–18]. Although there is genetic evidence for purifying selection in many species, including humans [10], it is unknown whether it occurs through the mechanisms identified in flies or if alternative mechanisms for purging mutant germline mtDNAs exist.

## Results

### PGCs eliminate mitochondria through intercellular cannibalism

To identify additional mechanisms of germline mtDNA control, we investigated how mtDNA quantity and quality are regulated in *C. elegans* PGCs. The entire *C. elegans* germ line descends from two PGCs, which remain quiescent during embryogenesis [19]. Although embryonic PGCs do not divide, they undergo a non-mitotic cellular remodeling process, discarding much of their cell mass and content. PGC remodeling occurs when PGCs form organelle-filled lobe-like protrusions, which adjacent endodermal cells cannibalize and digest (Fig. 1a) [20, 21]. Previously, we showed that PGCs lose much of their mitochondrial mass in the process of lobe cannibalism, suggesting that one role for this remodeling event could be to eliminate PGC mitochondria in bulk [20]. As such, lobe cannibalism might provide a novel mechanism for PGCs to adjust their mtDNA quantity and/or quality.

**Figure 1.**
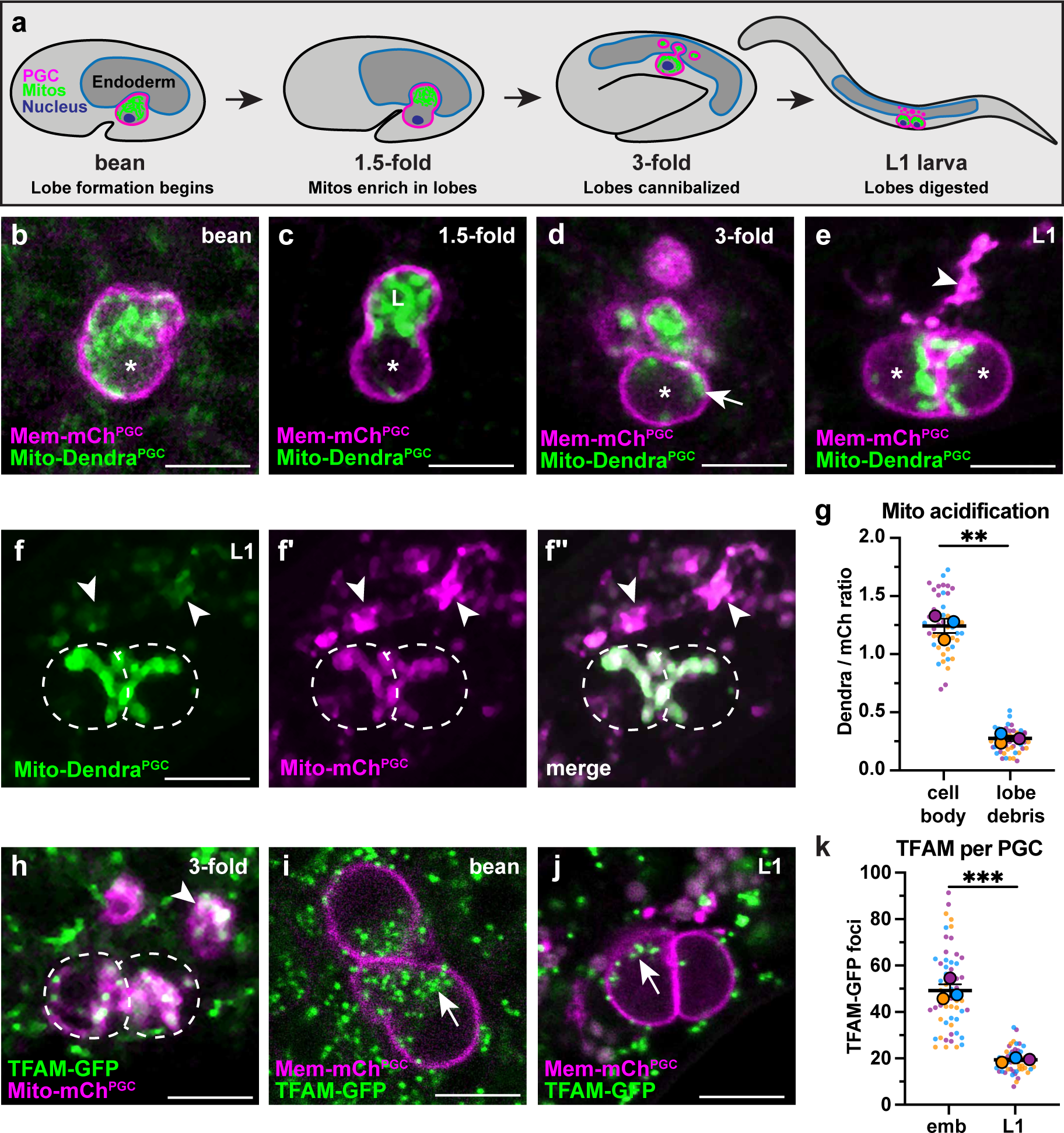
PGC lobe mitochondria and mtDNAs are cannibalized and digested (**a**). Schematic of PGC lobe formation and cannibalism. Bean stage to 3-fold embryos, one PGC visible; L1 larva, both PGCs visible. PGCs (magenta), PGC mitochondria (green) and endoderm (blue) are shown. (**b-e**) Plasma membranes and mitochondria in an embryonic PGCs just as lobes form (b), in PGCs with lobes (c-d) and in two L1 larval PGCs after lobes are digested (e; arrowhead, lobe debris in endoderm). *, nucleus, ‘L’, lobe. (**f-f”**) Acidified mitochondria (arrowheads) in digested PGC lobes of L1 larvae. Dashed lines, outline of PGC cell bodies. (**g**) Quantification of Mito-Dendra^PGC^ over Mito-mCh^PGC^ ratio in L1 PGCs revealing acidification in lobe debris relative to the cell body. **(h)** TFAM-GFP puncta within PGC mitochondria, present in both the cell body and in recently cannibalized lobes (arrowhead). Dashed lines, outline of PGC cell bodies. (**i-j**) TFAM-GFP in embryonic (i) and L1 larval (j) PGCs. (**k**) Quantification of TFAM-GFP foci in embryonic and L1 PGCs. Data in graphs are shown as a Superplot, with individual data points from three independent color-coded biological replicates shown as small dots, the mean from each experiment shown as a larger circle, the mean of means as a horizonal line, and the S.E.M as error bars.\*\**p* ≤ 0.01, \*\*\**p* ≤ 0.001, unpaired two-tailed Student’s *t*-test (k) and paired-ratio Student’s t-test (g). Scale bars, 5µm.

To begin to test this hypothesis, we used PGC-specific markers of the plasma membrane (PHPLC1∂1::mCherry, ‘Mem-mCh^PGC^’) and mitochondrial outer membrane (TOMM- 20^1-54^::Dendra2, ‘Mito-Dendra^PGC^’) to follow the distribution of PGC mitochondria during lobe formation and cannibalization in living embryos. Most PGC mitochondria moved into lobes shortly after they formed (Fig. 1b,c), but a subset returned to the cell body prior to lobe digestion (Fig 1d, S1a-c). Cell body mitochondria that are retained in L1 PGCs (Fig. 1e) represent the founding population present at the onset of larval germline expansion.

PGC lobe fragments present within endodermal cells co-localize with the lysosomal marker LAMP-1, suggesting that mitochondria within lobes are targeted for destruction and digested [20]. To test this hypothesis more directly, we labeled the mitochondrial outer membrane with Dendra2 (‘Mito-Dendra^PGC^’), which is pH-sensitive [22] and should be quenched when mitochondria are present within lysosomes. In L1 larvae, Mito-Dendra^PGC^ fluorescence was greatly reduced in cannibalized lobe mitochondria compared to pH- insensitive Mito-mCh^PGC^ [23] (arrowheads, Fig. 1f-g), whereas both markers labeled PGC cell body mitochondria robustly (dashed outline, Fig. 1f-g). We conclude that PGC lobe mitochondria are digested by endodermal cells shortly after lobes are cannibalized, permanently removing them from the mitochondrial pool passed on to L1 larval PGCs.

### Lobe cannibalism halves the number of PGC mtDNAs

To determine how elimination of mitochondria by lobe cannibalism affects the pool of germline mtDNAs, we first examined PGC mtDNAs visually. Mitochondrial transcription factor- A (TFAM), a component of the mtDNA nucleoid, is a well-characterized marker of mtDNA [24–26]. In human cells, individual TFAM nucleoids appear as puncta within the mitochondrial matrix and contain single, or at most a few, mtDNA genomes [27, 28]. We tagged the *C. elegans TFAM* homolog (*hmg-5*) endogenously with *gfp*. TFAM-GFP protein was expressed ubiquitously and formed puncta that localized to mitochondria, consistent with its known binding to mtDNA in *C. elegans* (Fig. S2a-b) [29]. Within PGCs, TFAM-GFP puncta were present in both cell body and lobe mitochondria, including those that had been recently cannibalized (arrowhead, Fig. 1h). The number of TFAM-GFP foci decreased more than two- fold between embryogenesis and the L1 larval stage (Fig. 1i-k), suggesting that lobe cannibalism results in a substantial loss of PGC mtDNAs.

To quantify the number of mtDNAs within PGCs, we developed a fluorescence activated cell sorting (FACS) protocol to purify GFP-labeled PGCs from either dissociated embryos or L1 larvae, which we paired with droplet digital PCR (ddPCR) to count mtDNA molecules per cell (Figs. 2a, S3a-d,g-h). We were able to isolate nearly pure populations of PGCs at both stages as determined by live imaging (Fig. S3e) and post-sort analysis (embryo: 98.0% +/- 0.5 pure; L1: 97.5% +/- 2.7 pure). Additionally, PGCs isolated from L1 larvae were less than half the volume of embryonic PGCs (Fig. S3e-f), indicating that lobe cannibalism had not yet initiated in most of the sorted embryonic PGCs and was complete in L1 PGCs, as expected [20].

**Figure 2.**
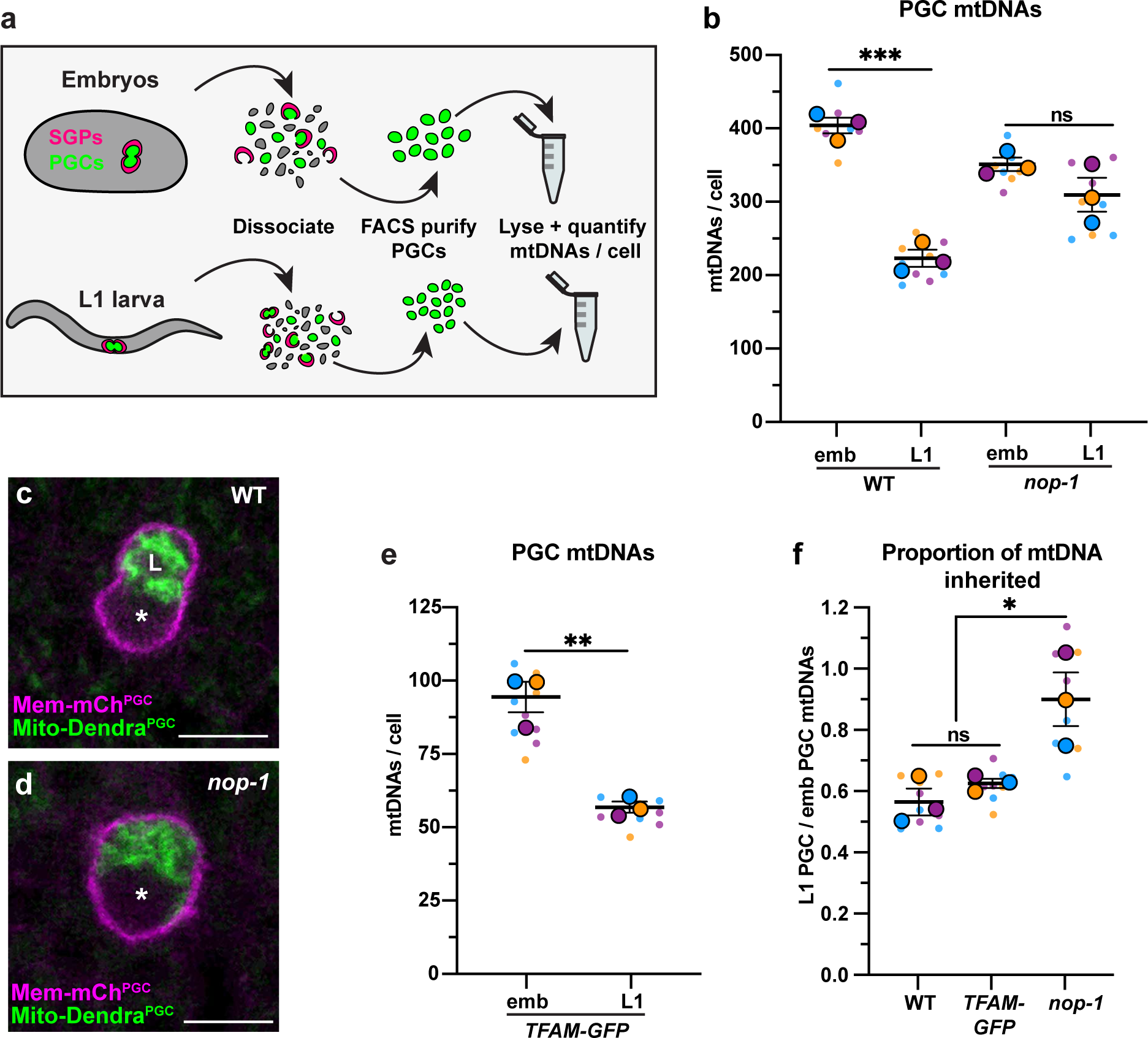
PGC lobe cannibalism eliminates a fixed fraction of mtDNAs (**a**) Schematic of FACS strategy to isolate PGCs from dissociated embryos and L1 larvae and quantify mtDNAs (see also Figure S3). (**b**) Quantification of mtDNA copy number per PGC in embryos and L1 of WT and *nop-1* mutants. (**c-d**) Mitochondria and plasma membrane in wild- type and *nop-1* mutant PGCs. (**e**) Quantification of mtDNA copy number per PGC in *TFAM- GFP* embryos and L1 larvae. (**f**) Proportion of embryonic PGC mtDNAs inherited by L1 PGCs in wild type, *TFAM-GFP,* and *nop-1* mutants (from data in b,e). Data in graphs: small dots are three technical replicates of ddPCR quantification from each of three color-coded biological replicates; the technical replicate mean from each experiment is shown as a larger circle, the mean of means as a horizonal line, and the S.E.M as error bars. n.s., not significant (*p*> 0.05), \**p* ≤ 0.05, \*\**p* ≤ 0.01, \*\*\**p* ≤ 0.001, unpaired two-tailed Student’s *t*-test. Scale bars, 5µm.

We determined that each embryonic PGC contained 401 +/- 11 mtDNAs (Fig. 2b), which is 1.2% of the number of mtDNAs we detected in whole early embryos (33,875 +/- 1819) (Fig. S2c). The volume of each PGC *in vivo* (275 +/- 7.5 µm^3^) is 1.2% of the volume of whole embryos (23,949 +/- 175 µm^3^), suggesting that PGCs inherit their mtDNAs from the pool present at fertilization through reductive embryonic cell divisions and that little or no mtDNA replication occurs prior to their birth. By contrast, L1 larval PGCs contained only 220 +/- 12 mtDNAs (Fig. 2b). The presence of fewer mtDNAs in L1 PGCs is consistent with our TFAM- GFP observations and suggests that PGC lobe cannibalism could eliminate nearly half of the mtDNA molecules that each PGC inherits at its birth (Fig. 2f).

To directly test whether lobe cannibalism causes the loss of mtDNAs that occurs within PGCs between embryonic and L1 stages, we examined PGC mtDNA number in *nop-1* mutants, in which most PGCs fail to form lobes (Fig. 2c,d) [30]. *nop-1* mutant L1 PGCs retained a significantly higher proportion of embryonic PGC mtDNAs compared to wild type (Fig. 2b,f). This finding implicates lobe cannibalism in the two-fold reduction in mtDNA that occurs as PGCs transition from embryogenesis to the L1 stage.

Lobe cannibalism could reduce the number of mtDNAs to a fixed number, or alternatively, eliminate a fixed proportion of the mtDNAs present within PGCs regardless of how many are present. To distinguish between these possibilities, we took advantage of the fact that changing TFAM activity can alter mtDNA copy number [31, 32]. Indeed, we found that whole embryos from the *TFAM-GFP* knock-in strain contained significantly fewer mtDNAs (8630 +/- 662) than wild type (Fig. S2c), indicating that the GFP tag partially interferes with TFAM function. Using the *TFAM-GFP* strain, we asked how many mtDNAs PGCs eliminate during lobe cannibalism if they are born with a reduced number. *TFAM-GFP* embryonic PGCs contained 94 +/- 5 mtDNAs (Fig. 2e) and *TFAM-GFP* L1 PGCs contained 56 +/- 2 mtDNAs (Fig. 2e). Thus, despite the presence of markedly fewer mtDNAs in the *TFAM-GFP* strain, L1 PGCs still inherit a comparable percentage of the mtDNAs contained within embryonic PGCs (wild-type: 55%, *TFAM-GFP*: 60%) (Fig. 2f). These data indicate that lobe cannibalism does not subtract the number of PGC mtDNAs to a defined number, but rather divides the population of PGC mtDNAs present by a fixed proportion.

### Lobe cannibalism creates a bottleneck and establishes an mtDNA set point in germline stem cells

Our results so far suggest that lobe cannibalism could create a germline mtDNA bottleneck by halving the number of PGC mtDNAs. However, if the initial cycles of larval germline proliferation proceed in the absence of mtDNA replication, the number of mtDNAs per germ cell would continue to drop and the bottleneck would occur at a later stage of germline development. When L1 larvae first encounter food, PGCs exit from quiescence and begin to proliferate, forming a population of undifferentiated germline stem cells (GSCs) [19, 33]. It is not known whether germline mtDNA replication has begun at this stage. Previous qPCR experiments on whole worms first revealed a significant expansion of germline mtDNAs after the L3 larval stage [34, 35]. However, these experiments may have lacked the resolution to detect an increase in mtDNAs were it to occur within the relatively small number of GSCs present in whole L1 larvae.

To determine if mtDNAs replicate as L1 PGCs exit quiescence and divide to produce GSCs, we quantified TFAM foci as PGCs in fed L1s began to proliferate as GSCs. To circumvent the mtDNA replication defects that we noted in *TFAM-GFP* worms, we tagged *TFAM* (*hmg-5*) endogenously with a much smaller tag, *gfp(11)* [36, 37]. To visualize TFAM-GFP(11), we expressed a mitochondrially targeted, PGC-specific GFP(1-10) [Mito-GFP(1-10)^PGC^]. GFP(1-10) alone was minimally fluorescent, but upon binding to GFP(11) formed a functional fluorophore (Fig. S4a,b). Within PGCs, TFAM-GFP(11) showed an identical localization pattern to TFAM-GFP in PGCs (Fig. 3a,b), and did not cause significant defects in mtDNA copy number (Fig. S4c). Larvae fed beginning at the L1 stage showed a progressive increase in the number of TFAM-GFP(11) foci per germ line (Fig. 3c-f, compare to 3b). TFAM- GFP(11) foci numbers began to increase even before the first division of the PGCs was complete (early L1, Fig. 3c,g), and continued to expand through the L2 stage (an average of 22 GSCs), when we stopped our analysis (Fig. 3f,g). In contrast to the increasing number of TFAM-GFP(11) foci per *germ line* over this period, the number of foci per *germ cell* remained constant after a transient spike in early L1s (2 cells), and stabilized at a number of foci similar to that of L1 that had not been fed (Fig. 3h).

**Figure 3.**
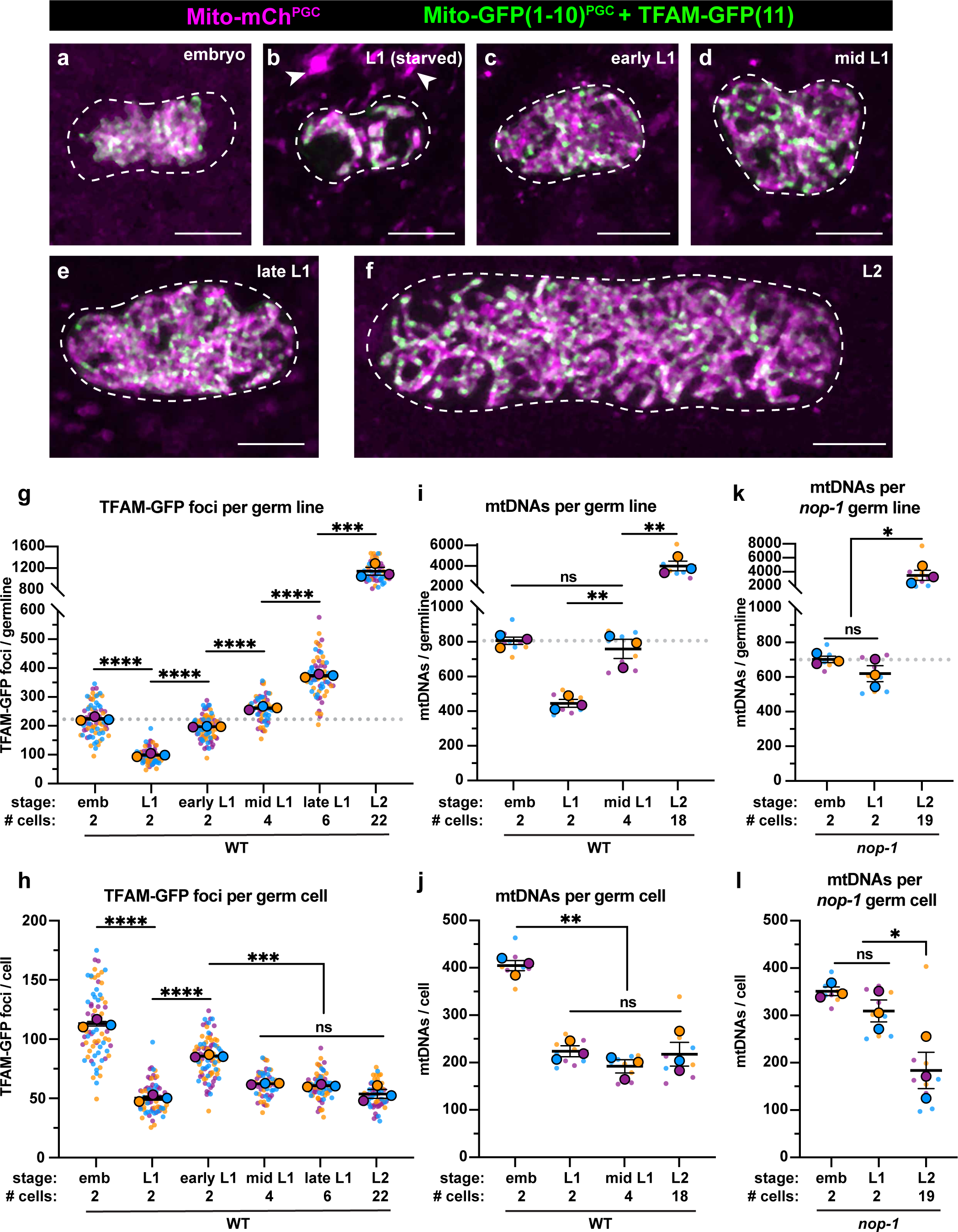
PGC lobe cannibalism generates an mtDNA bottleneck and set point (**a-f**) Germline mitochondria and TFAM-GFP(11) in live embryos and larvae at the indicated stage. Dashed lines outline the PGCs or GSCs. (**g-h**) Quantification of TFAM-GFP(11) foci per germ line (g) and per germ cell (h) in embryos and larvae. (**i-j**) Quantification of mtDNAs per germ line (i) or per germ cell (j) in embryos and larvae; data shown for PGC mtDNA copy number in embryos and starved L1s are provided for comparison and originate from Fig. 2 panel (b). (**k-l**) Quantification of mtDNAs per germ line (k) or per germ cell (l) in *nop-1* mutant embryos and larvae; data shown for PGC mtDNA copy number in *nop-1* mutant embryos and starved L1s are provided for comparison and originate from Fig. 2 panel (b). Data in graphs: small dots are individual animals [TFAM-GFP(11) measurements] or technical replicates (ddPCR experiments) from three color-coded biological replicates; the mean from each experiment is shown as a larger circle, the mean of means as a horizonal line, and the S.E.M as error bars. n.s., not significant (*p*> 0.05), \**p* ≤ 0.05, \*\**p* ≤ 0.01, \*\*\**p* ≤ 0.001, \*\*\*\**p* ≤ 0.0001 unpaired two-tailed Student’s *t*-test. Scale bars, 5µm.

To complement these experiments, we sorted GSCs (Fig. S5) from fed L1 larvae (containing an average of 4 GSCs) and L2 larvae (containing an average of 18 GSCs), and counted the number of mtDNA molecules per GSC. Consistent with our TFAM-GFP(11) observations, the number of mtDNAs per germ line increased over this period (Fig. 3i), although the number of mtDNAs per GSC remained constant, and similar to that in starved L1s (∼200) (Fig. 3j). Together, these results indicate that mtDNAs replicate as L1 PGCs begin to divide to form GSCs, and thereafter balance mtDNA replication with cell division to maintain a constant number of mtDNAs per GSC, through at least the L2 stage.

The observation that GSCs contain the same number of mtDNAs as L1 PGCs suggests that lobe cannibalism might function to reduce PGC mtDNA numbers to an optimal level. To explore this hypothesis, we examined mtDNA number in GSCs of *nop-1* mutants, since we found that *nop-1* L1 PGCs contain excess mtDNAs. Remarkably, *nop-1* GSCs isolated from L2 larvae contained a similar number of mtDNAs (182 +/- 38) as did wild-type GSCs (215 +/- 25) (Fig. 3k,l). These results suggest that GSCs actively regulate mtDNA replication to maintain ∼200 mtDNAs per cell, even if excess mtDNAs are present at the onset of germline expansion.

### Purifying selection reduces mutant mtDNA heteroplasmy in PGCs independently of lobe cannibalism

Our experiments so far have not addressed whether lobe cannibalism eliminates mitochondria indiscriminately, or alternatively, if poorly functioning mitochondria, containing high levels of mutant mtDNA, are preferentially targeted to lobes for destruction. To examine this question, we investigated PGCs containing the *uaDf5* mtDNA deletion. *uaDf5* removes 3.1 kb of the mitochondrial genome, including several essential genes (Fig. 4a), and therefore can exist only when in heteroplasmy with wild-type mtDNA [38]. However, *uaDf5* persists stably because it is preferentially replicated compared to wild-type mtDNA ([29, 39–41]. Our experiments above suggest that there is little or no mtDNA replication occurring in PGCs, potentially providing an opportunity for purifying selection to reduce *uaDf5* levels before larval germline expansion begins.

**Figure 4.**
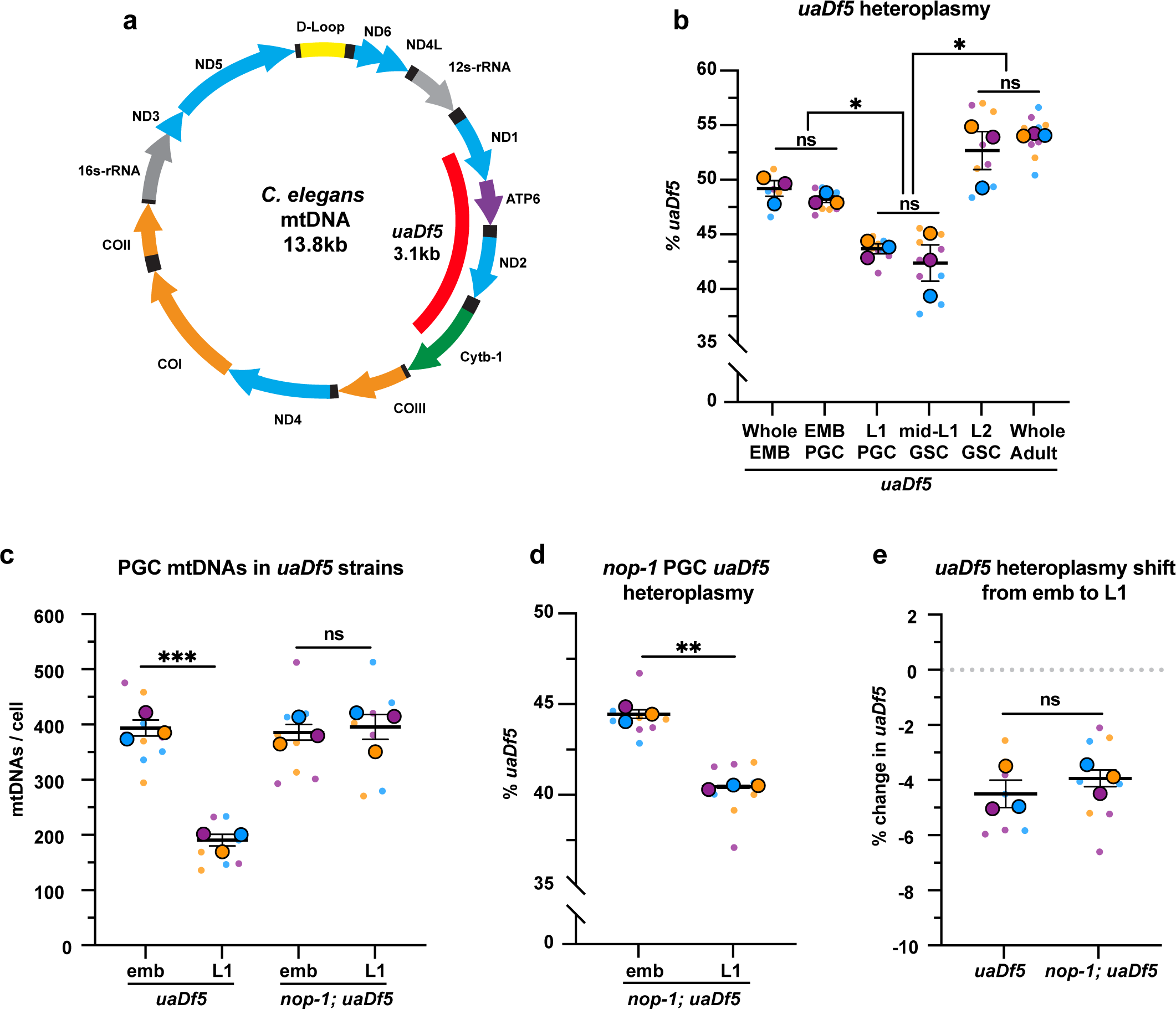
PGCs reduce *uaDf5* heteroplasmy independently of lobe cannibalism (a) Schematic of *C. elegans* mtDNA; genes are indicated with colored arrows and the region deleted in *uaDf5* is shown with a red bar. (**b**) Quantification of *uaDf5* heteroplasmy in whole embryos, sorted PGCs or GSCs, or whole adults at the indicated stages. (**c**) Quantification of mtDNA copy number in PGCs of *uaDf5* and *nop-1; uaDf5* mutants. (**d**) Quantification of *uaDf5* heteroplasmy in *nop-1; uaDf5* mutant PGCs. (**e**) Data from (b and d) presented as change in heteroplasmy shift from embryonic to L1 PGCs. Data in graphs: small dots are three technical replicates of ddPCR quantification from each of three color-coded biological replicates; the technical replicate mean from each experiment is shown as a larger circle, the mean of means as a horizonal line, and the S.E.M as error bars. n.s., not significant (*p*> 0.05), \**p* ≤ 0.05, \*\**p* ≤ 0.01, \*\*\**p* ≤ 0.001, paired (b, d) and unpaired (b, c, e) two-tailed Student’s *t*-test.

First, we measured *uaDf5* heteroplasmy in whole embryos, embryonic PGCs, and L1 PGCs, as well as in GSCs of fed larvae (Fig. 4b, S6). *uaDf5* occurred at 48% +/- 0.3 heteroplasmy in embryonic PGCs, which was nearly identical to its heteroplasmy in whole embryos (Fig. 4b), suggesting that there is not strong selection against *uaDf5* during embryogenesis prior to PGC birth. However, in L1 PGCs, *uaDf5* heteroplasmy dropped by 4.5% (Fig. 4b). This effect was not specific to *uaDf5*, as PGC heteroplasmy of the 1.5kb *mptDf2* deletion was also reduced between embryogenesis and the L1 stage (Fig. S6). Within starved L1 PGCs, *uaDf5* was present at 43.5% +/- 0.5 heteroplasmy, and was maintained at a similar level after the PGCs divided once to form 4 GSCs (mid-L1 stage). However, by the L2 stage (average of 20 GSCs), *uaDf5* heteroplasmy increased to 53% +/- 1.7 - a level nearly identical to that of whole adult worms (Fig. 4b). These findings suggest that PGCs utilize purifying selection to reduce levels of mutant mtDNAs at a stage when they cannot take advantage of mtDNA replication to expand selfishly within the germ line. However, once mtDNA replication begins in larval GSCs, the frequency of mutant mtDNAs can once again rise.

To test whether lobe cannibalism is responsible for purifying selection against *uaDf5* in PGCs, we examined *uaDf5* heteroplasmy in *nop-1* mutant PGCs. Similar to wild type, *uaDf5* PGCs reduced their total mtDNA content ∼two-fold as a result of lobe removal, and as expected, *nop-1; uaDf5* PGCs failed to reduce their mtDNA (Fig. 4c). Surprisingly, we found that in *nop-1; uaDf5* mutants, *uaDf5* heteroplasmy still decreased from embryonic PGCs to L1 PGCs (Fig. 4d,e). We conclude that lobe cannibalism is not responsible for the reduction in *uaDf5* heteroplasmy within PGCs, implicating an alternative pathway in PGC mtDNA purifying selection.

### Autophagy eliminates a subset of PGC mitochondria non-selectively

Cells can use autophagy to remove damaged cellular components and organelles, including mitochondria. During autophagy, an autophagosome membrane encapsulates organelles and cytoplasm, subsequently fusing with a lysosome to degrade its contents [42]. To address whether mtDNA purifying selection in PGCs might be mediated by autophagy, we first used the pH-discriminating Mito-mCh^PGC^ and Mito-Dendra^PGC^ reporters to observe whether any PGC mitochondria become acidified before lobe cannibalism occurs. In the majority of *uaDf5* embryos, we observed one or more large, distinct foci of acidified mitochondria [mCherry(+) Dendra(-)] within PGCs prior to lobe cannibalism (Fig. 5a,c).

**Figure 5.**
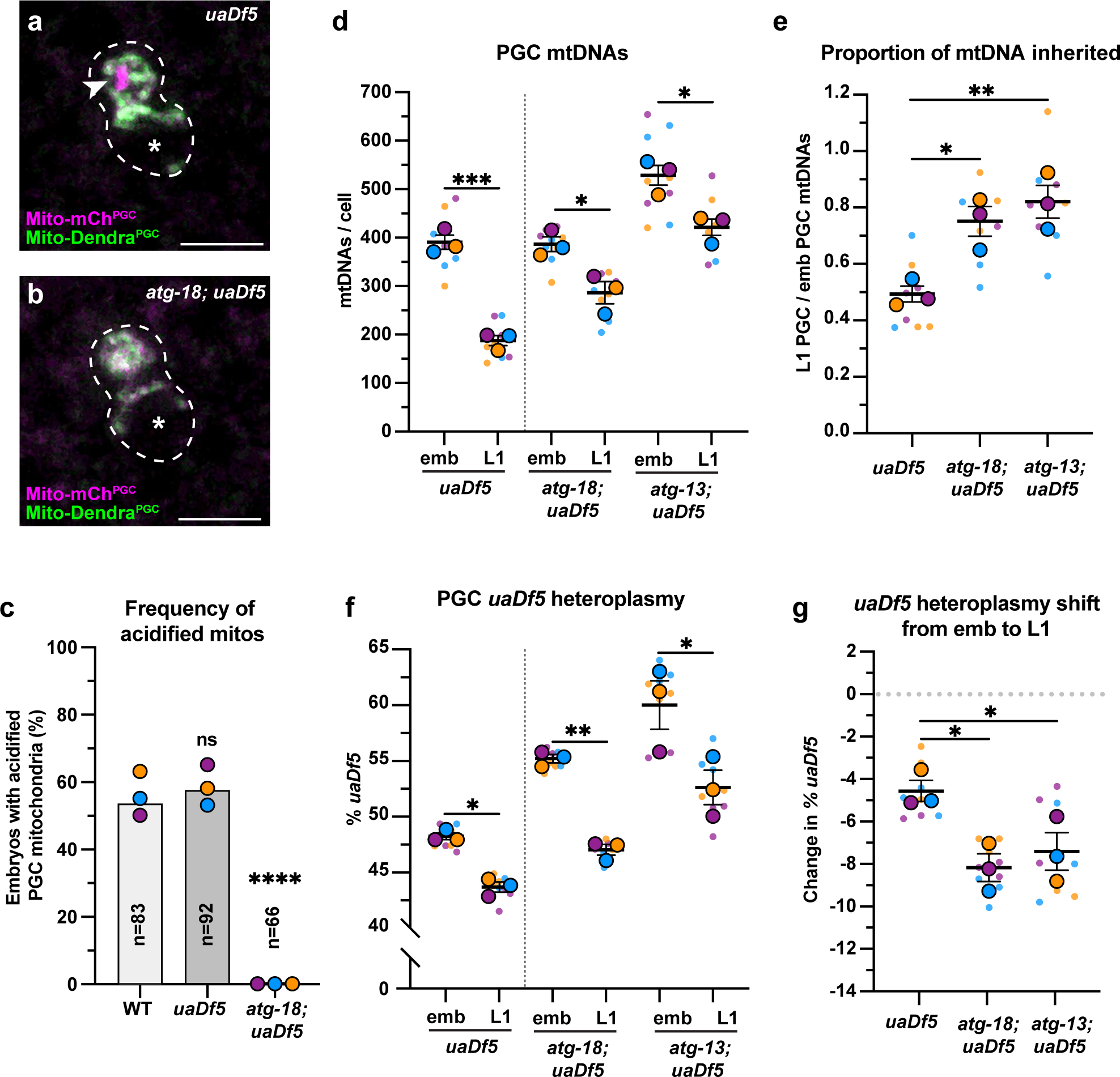
Autophagy eliminates a pool of PGC mtDNAs non-selectively (**a-b**) Acidified mitochondria (red regions, arrowhead in a) in *uaDf5* PGCs (a) and absent in *atg-18; uaDf5* PGCs. (**c**) Percent of embryos with acidified mitochondria in PGCs. Three biological replicates (N ≥ 16) are shown as colored circles, with peak of the bar on the graph representing the mean. Fisher’s exact test was used to determine statistical significance. n.s., not significant (*p*> 0.05), **** *p* ≤ 0.0001 (**d**) mtDNA copy number in *atg-18*; *uaDf5,* and *atg-13; uaDf5* embryonic and L1 PGCs; data shown for *uaDf5* are provided for comparison, originate from Figure 4 panel (c), and are delineated with a dashed line. (**e**) Data from (d) presented as proportion of embryonic PGC mtDNAs inherited by L1 PGCs. (**f**) *uaDf5* heteroplasmy in *atg-18*; *uaDf5,* and *atg-13; uaDf5* PGCs; data shown for *uaDf5* are provided for comparison, originate from Figure 4 panel (b), and are delineated with a dashed line. (**g**) Data from (f) presented as change in heteroplasmy shift from embryonic to L1 PGCs. Data in graphs: small dots are three technical replicates of ddPCR quantification from each of three color-coded biological replicates; the technical replicate mean from each experiment is shown as a larger circle, the mean of means as a horizonal line, and the S.E.M as error bars. n.s., not significant (*p*> 0.05), \**p* ≤ 0.05, \*\**p* ≤ 0.01, \*\*\**p* ≤ 0.001 paired (f) and unpaired (d, e, g) two-tailed Student’s *t*-test.

Acidified foci were completely absent in *atg-18/WIPI2; uaDf5* null mutant embryos (Fig. 5b,c), in which autophagy is blocked at the autophagosome membrane nucleation step [43], suggesting that the acidified foci are PGC mitochondria that are eliminated by autophagy.

To test whether autophagy preferentially removes *uaDf5* mtDNAs in PGCs, we sorted embryonic and L1 PGCs in *uaDf5* mutants with putative null mutations in *atg-18* and *atg-13*, which blocks autophagy at an earlier initiation step [43, 44]. L1 PGCs in *atg-18* and *atg-13* mutants had reduced numbers of total mtDNAs compared to embryonic PGCs, although a smaller percentage (*atg-18:* 25%, *atg-13:* 20%) of mtDNAs were eliminated compared to *uaDf5* alone (52%) (Fig. 5d,e). These findings suggest that the autophagy pathway acts in parallel with lobe cannibalism and is partially responsible for the reduction of mtDNAs in PGCs.

Unexpectedly, in both *atg-13; uaDf5* and *atg-18; uaDf5* mutants, *uaDf5* heteroplasmy was still reduced in L1 PGCs compared to embryonic PGCs (Fig. 5f,g). We conclude that autophagy likely eliminates a subset of mitochondria and mtDNAs within PGCs non-selectively, but is not responsible for purifying selection against *uaDf5* mutant mtDNA. Consistent with this interpretation, wild-type embryonic PGCs contained foci of acidified mitochondria at comparable frequencies to *uaDf5* PGCs (Figs. 5c, S7).

### PINK1 mediates autophagy-independent mtDNA purifying selection in PGCs

The PINK1/Parkin signaling pathway, which consists of the mitochondrial kinase PINK1 and its effector ubiquitin ligase Parkin, can recognize and mark defective mitochondria for destruction either via autophagy, or through autophagy-independent pathways [16, 45, 46]. To address whether homologs of PINK1 or Parkin are required for purifying selection of *uaDf5*, we examined *uaDf5* heteroplasmy in PGCs with putative null mutations in *pink-1/PINK1*, *pdr- 1/Parkin,* and double mutants. As expected, single and double mutants had reduced mtDNA content in L1 PGCs compared to embryonic PGCs, although to a lesser extent than *uaDf5* controls (Fig. 6a,b). However, even though *uaDf5* heteroplasmy was markedly higher in all three backgrounds compared to *uaDf5* controls (see Discussion), only *pink-1,* and *pink-1; pdr- 1* double mutants, but not *pdr-1* single mutants, abrogated the reduction in *uaDf5* heteroplasmy between embryonic and L1 PGCs (Fig. 6c-d). Taken together, these findings indicate that PINK1 alone, acting independently of Parkin, is required for autophagy- independent purifying selection against mutant mtDNAs within *C. elegans* PGCs.

**Figure 6.**
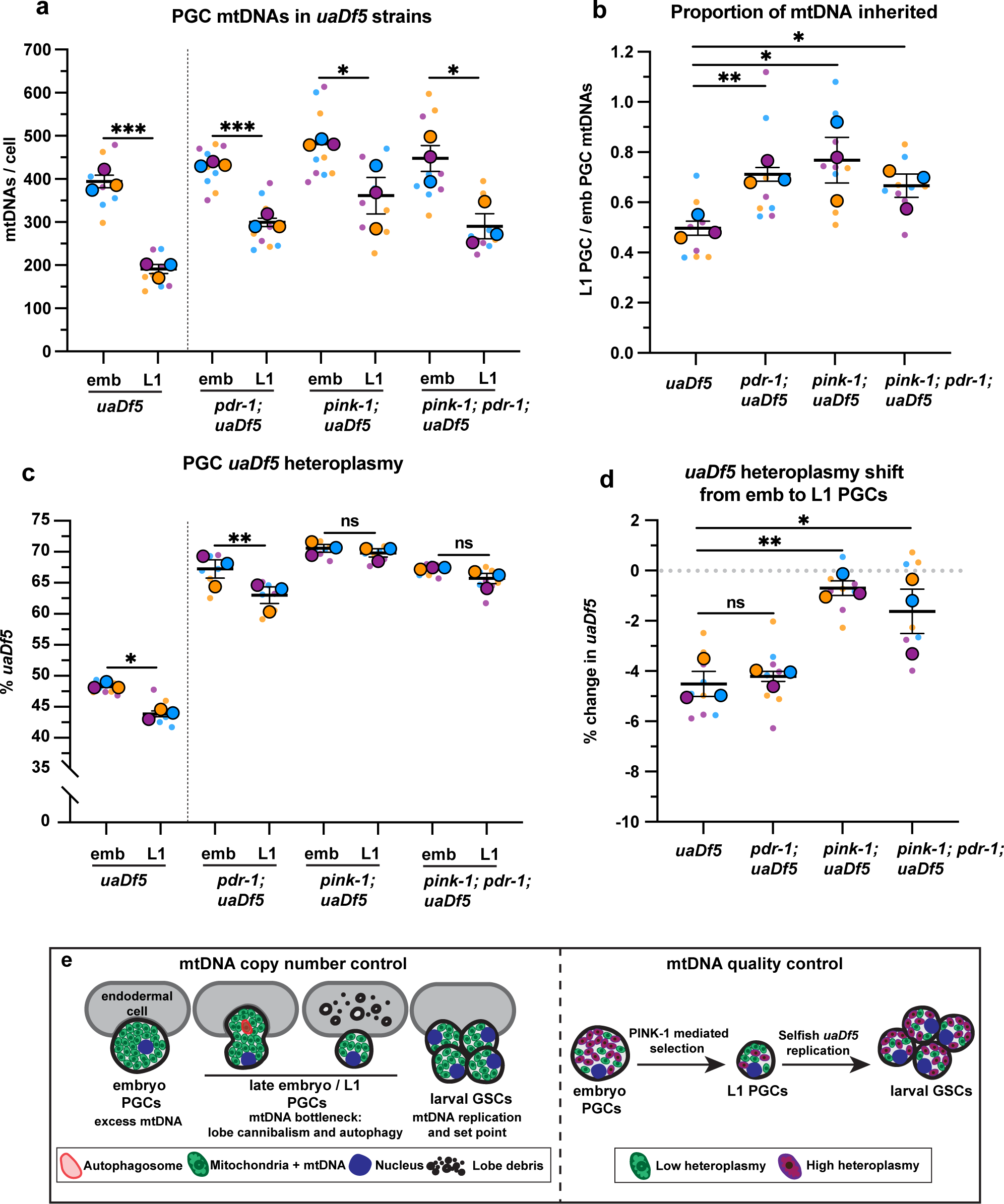
The kinase PINK-1 mediates mtDNA purifying selection in PGCs (**a**) mtDNA copy number in *pdr-1; uaDf5, pink-1; uaDf5,* and *pink-1; pdr-1; uaDf5* embryonic and L1 PGCs; data shown for *uaDf5* are provided for comparison, originate from Figure 4 panel (c), and are delineated with a dashed line. (**b**) Data from (a) presented as proportion of embryonic PGC mtDNAs inherited by L1 PGCs. (**c**) Percent *uaDf5* heteroplasmy in *pdr-1; uaDf5, pink-1; uaDf5,* and *pink-1; pdr-1; uaDf5* ; data shown for *uaDf5* are provided for comparison, originate from Figure 4 panel (b), and are delineated with a dashed line. Data in graphs: small dots are three technical replicates of ddPCR quantification from each of three color-coded biological replicates; the technical replicate mean from each experiment is shown as a larger circle, the mean of means as a horizonal line, and the S.E.M as error bars. n.s., not significant (*p*> 0.05), \**p* ≤ 0.05, \*\**p* ≤ 0.01, \*\*\**p* ≤ 0.001, paired (c) and unpaired (a, b, d) two- tailed Student’s *t*-test. (**e**) Model for regulation of mtDNA quantity and quality in PGCs and GSCs.

## Discussion

Our findings show that *C. elegans* PGCs actively regulate both mtDNA quantity and quality, but do so through independent and parallel mechanisms (Fig. 6e). The cannibalism of PGC lobes creates a bottleneck and mtDNA setpoint, whereas PINK1 preferentially reduces mutant mtDNA heteroplasmy. We propose that this combined regulation optimizes the founding population of mitochondria before expansion and differentiation of the germ line.

Elimination of mitochondria through intercellular cannibalism provides a previously undescribed mechanism for bottleneck formation. We postulate that embryonic PGCs inherit excess maternal mtDNAs, as they are born from relatively few embryonic cell divisions [21], and lobe cannibalism halves PGC mtDNA copy number to establish a level that is maintained in GSCs as mtDNA replication ensues. Having ∼200 mtDNAs per PGC appears to be important, as even when L1 PGCs inherit an excess of mtDNAs in *nop-1* mutants, mtDNA copy number quickly resets to ∼200 shortly after PGCs differentiate into proliferating GSCs.

These findings indicate that GSCs balance mtDNA replication with cell division to reach an mtDNA set point of ∼200 that is actively maintained. It is possible that this number of mtDNAs is needed for sufficient selection against deleterious mtDNA mutations, minimizes damaging free radical production [20], or is optimal for balancing mitochondrial function with germ cell size and physiology.

Whereas PGC lobe cannibalism produces a stochastic reduction in mtDNA number, we found that PINK1 specifically reduces the fraction of mutant mtDNAs in PGCs. While the effect of PINK1-mediated selection against *uaDf5* is moderate, even small decreases in heteroplasmy could potentially have important evolutionary consequences. For example, individual selection events against *de novo* mtDNA mutations could eliminate them from the germ line permanently. In other systems, PINK1 can eliminate poorly functioning mitochondria by recruiting Parkin and inducing mitophagy [47]. However, we find no role for Parkin or autophagy in PGC mtDNA purifying selection, although non-selective autophagy is partially responsible for reducing mtDNA number. Alternative mechanisms of mitochondrial elimination have been described in cultured mammalian cells, such as direct targeting to endolysosomes [45, 46]. It will be important in future studies to determine whether PINK1 operates in PGCs through one of these pathways or via a novel mechanism. It is worth noting that *uaDf5* heteroplasmy in embryonic PGCs is higher in *pink-1*, *pdr-1*, and autophagy mutants, suggesting that these pathways have roles in purifying selection during other stages of germ line development, as they do in somatic cells [48–50].

Purifying selection in *C. elegans* PGCs differs from mechanisms that operate in the *Drosophila* ovary, where mutant mtDNAs are eliminated through mitochondrial fission followed by autophagy, and mutant mtDNA replication is selectively inhibited by PINK1 [14–18].

Although it is difficult to exclude the possibility that very small amounts of mtDNA replication occur in *C. elegans* PGCs, we do not detect robust mtDNA replication until PGCs differentiate into GSCs. This finding is also supported by our observation that *uaDf5* heteroplasmy decreases in PGCs, whereas *uaDf5* is known to selfishly expand through preferential mtDNA replication [29]. Indeed, we showed that as PGCs differentiate to GSCs, *uaDf5* heteroplasmy rapidly increases to levels found in the adult. Previous work has suggested that selection also occurs during *C. elegans* oogenesis, although the mechanism is unknown [40, 51]. It will be interesting to determine if these different means of achieving purifying selection are stage- specific (ovary versus PGC), or reveal that multiple mechanisms can be used toward the common goal of eliminating mutant mtDNA genomes from the germ line.

## Acknowledgements

We thank the Caenorhabditis Genetics Center (CGC), Heng-Chi Lee (U. of Chicago) and Dustin Updike (MDI Biological Laboratory) for providing worm strains. The CGC is supported by the NIH Office of Research Infrastructure Programs (P40 OD010440). We thank members of the Nance laboratory, Florenal Joseph, and Melissa Pamula for comments on the manuscript. We thank Peter Lopez, James Alvarado, Yulia Chupalova, and Sitharam Ramaswami for FACS/ddPCR assay development, and Michael Cammer for help with image analysis and acquisition. FACS was performed at the NYULMC Cytometry and Cell Sorting Laboratory; ddPCR was performed at the NYULMC Genome Technology Center; and Microscopy used instrumentation in the NYULMC Microscopy Laboratory, all of which are partially supported by the Laura and Isaac Perlmutter Cancer Center support grant P30CA016087 from the National Institutes of Health/National Cancer Institute. This work was supported by fellowships/training grants from the National Institutes of Health and NYSTEM to AZAS (NIH: F31HD102161, NYSTEM: C32560GG) and a research grant from the National Institutes of Health to JN (R35GM118081).

## Author contributions

AZAS and JN conceived the project. AZAS performed all experiments except creation of the Mito-Dendra^PGC^ marker, which was made by YA. AZAS and JN analyzed and interpreted the data. NT and MP provided training in cell sorting and identified the *mptDf2* strain. AZAS and JN wrote the initial manuscript, and all authors edited the manuscript.

## Methods

### Worm culture and strains

Unless otherwise stated, all strains were maintained at 20°C on nematode growth medium plates seeded with *Escherichia coli* OP50 according to standard methods [52]. For egg isolation and L1 synchronization, semi-synchronized L1 larvae were outgrown on 10cm Enriched Peptone plates seeded with *E. coli* NA22. Gravid adults were then washed off and early stage embryos were isolated via worm bleaching. Isolated eggs were broken into two populations - one for immediate embryo dissociation and another which was allowed to hatch overnight in M9 for L1 synchronization/dissociation. For L1 feeding experiments, synchronized L1 larvae were plated onto peptone plates and grown for 12 and 24 hrs at 20°C (for cell sorting), or for 6, 9, 12, and 24 hours at 23°C (for live imaging). A list of all strains used/generated in the study is available in Supplementary Table 1.

### PGC isolation and cell sorting

Cell dissociation of early embryos and larvae was performed as described previously [53, 54] with slight modifications described in detail below.

#### Embryonic cell dissociation

Purified embryos were pelleted at 3000g for 30 seconds in non-stick 1.5mL tubes, resuspended in 600µL chitinase (Sigma C6317) (2mg/mL) in conditioned-egg buffer (25 mM HEPES (Sigma H3375) pH 7.3, 118 mM NaCl, 48mM KCl, 2 mM CaCl2, 2 mM MgCl2, adjusted to mOsm 340±5 with ddH2O), hereafter referred to as egg buffer, and incubated on a rocking nutator for 15 minutes at room temperature. After 15 minutes, 800µL of cold egg buffer was added, embryos were spun at 900g for 4 minutes at 4°C, and then resuspended in 800µL Accumax-egg buffer solution (Innovative Cell Technologies, AM105, 1:3 dilution ratio in egg buffer). For dissociation, embryos were pipetted up and down ∼80 times using a p1000 pipette. To wash away debris, dissociated embryos were spun at 900g for 4 minutes at 4°C a total of three times. Washed cells were resuspended in 800µL of cold egg buffer, and single cells were separated from clumps by gravity settling on ice for 15-20 minutes. For *uaDf5* heteroplasmy experiments, 25µL of dissociated cells was removed at this stage, mixed 1:1 with worm lysis buffer, lysed as described below, and stored at -80°C for ddPCR.

#### Larval cell dissociation

Dissociation of larvae was performed at three stages: starved L1s, mid-L1s (L1s fed 12 hrs, 20°C) and L2s (L1s fed 24hrs, 20°C). Larvae at a specific stage were collected into 15mL conical tubes, spun down at 3000g for 30 seconds, and washed with ddH2O 2-6X. Larvae were then collected into 1.5mL non-stick tubes and spun at 16,000g for 2 minutes. Depending on the size of the pellet, larvae were split into multiple tubes such that each tube had no more than 100µL of pelleted animals. Starved L1s, mid-L1s and L2s were then resuspended in 250µL of SDS-DTT solution (20mM HEPES pH 8.0, 0.25% SDS (Sigma 71725), 200mM DTT (Sigma D0632), 3% sucrose) and incubated 2, 2.5, and 3 minutes respectively with gentle mixing. To stop the reaction 1mL of cold egg buffer was added, then animals were spun at 16,000g for 1 minute and washed an additional 5X with cold egg buffer. Following the last wash, SDS-DTT treated animals were resuspended in 250µL pronase (Sigma P8811) solution (15mg/mL in egg buffer) and incubated for 5-15 minutes on a rocking nutator at room temperature. Animals were then dissociated by trituration with a pipet for an additional 25 minutes (∼60 times every 5 minutes) in pronase solution. To end the dissociation, 1mL of cold egg buffer was added and cells were spun down at 9600g for 3 minutes at 4°C. Cell pellets were resuspended in 1mL of cold egg buffer and washed 3X by spinning 1600g for 6 minutes at 4°C. Following the final wash, dissociated cells were resuspended in 800µL of cold egg buffer and separated from undissociated larvae and clumps by gravity settling on ice for 30-40 minutes.

#### FACS and PGC isolation

For sorting experiments, we used a strain expressing endogenously tagged GLH-1-GFP, which is a germline-specific protein [55], as well as a transgenic mCherry marker specific to somatic gonad precursor cells (SGPs) [56], which ensheath the PGCs and are the most likely contaminating population of cells. Approximately 15 minutes prior to cell sorting, DAPI (Sigma D9542) was added to the cells (final concentration of 0.125ug/mL) as a viability marker. GLH- 1-GFP(+) ; SGP-mCherry(-) ; DAPI (-) cells were isolated via FACS using a 100µm nozzle on a BD FACSAriaII cell sorter. For quality control, sorted cells were live imaged (see ‘microscopy’ below) to confirm the presence of GFP(+); RFP(-) cells. Purity was assayed, via post-sort analysis (N=3), by resorting cells and quantifying the percentage of GFP(+) ; RFP(-) ; DAPI(-) cells in the population in FlowJo software V10. For most ddPCR analysis, 1000-5000 PGCs were sorted into 500µL of 0.5X worm lysis buffer (recipe below) in a screw-cap 1.5mL microfuge tube (20,000 and 10,000 cells were sorted for wild-type and *TFAM-GFP* PGCs respectively). Following sorting, PGCs were lysed for 30 minutes on ice and then incubated in a table-top heating block for 1 hour at 55°C followed by 15 minutes at 95°C. Cell lysates were frozen at -80°C until needed for ddPCR. For live imaging, 1000-2500 PGCs were sorted into 500µL of conditioned L-15 medium (10% FBS, 50 U/mL penicillin + 50 μg/mL streptomycin (Sigma P4458), adjusted to mOsm 340±5 with 60% sucrose) and kept on ice. Embryonic and larval PGCs were spun down at 900g (4 minutes) and 1600g (6 minutes) respectively, all but 50µL of conditioned L-15 was removed, and cells were gently resuspended for imaging (see ‘microscopy’ below).

### Quantitative PCR (qPCR)

#### Whole L4 larvae mtDNA copy number

For standard curve generation, an 887bp portion of mtDNA containing *nd-1* was amplified by PCR and TA-cloned into pMiniT2.0 using the NEB PCR cloning kit (NEB E1202S). Purified plasmid was linearized with *Bam*HI-HF (NEB 3136), and DNA concentration was quantified using a Nanodrop spectrometer (Thermo scientific). For the standard curve, 64,000, 32,000, 24,000, 16,000, 12,000, 8000, 6000, and 4000 copies of plasmid were run in triplicate as described below. Oligos targeting the mitochondrial gene *nd-1*(see ‘Droplet digital PCR’ below) were used for qPCR quantification. For absolute quantification, single late-L4 larvae were picked into 5µL of worm lysis buffer (50 mM KCl, 10 mM Tris-HCl (pH 8.0), 2.5 mM MgCl2, 0.45% IGEPAL (Sigma I8896), 0.45% Tween 20 (Sigma P9416), 0.01% gelatin (Sigma G1393), and 200 µg/mL proteinase K (Invitrogen 2530049) and flash frozen at -80°C for 15 minutes. Worms were then lysed in a thermal cycler at 60°C for 1 hour followed by 15 minutes at 95°C. Prior to qPCR, lysed L4s were diluted 20X by adding 95µL of nuclease-free water (Invitrogen 4387936) and mixed thoroughly by pipetting. 8µL of lysate (or diluted plasmid for standard curve) was used in triplicate for each individual sample. qPCR was performed as a 20µL reaction with 500µM of each primer, using BioRad 2X SsoAdvanced Universal SYBR Green Supermix (BioRad 1725271) in a Roche LightCycler 480 machine. The PCR program was as follows: 10 minutes at 98°C, 40 cycles of 98°C for 15s and 60°C for 1 minute. Crossing point (Cp) values were derived using the Second Derivative Maximum method of the Roche LightCycler 480 software.

### Whole embryo lysis

Embryos were isolated from gravid adults and chitinase treated for 8 minutes at room temperature to dissolve the egg shell prior to lysis. Chitinase-treated embryos were washed 2- 3X with cold egg buffer and transferred to a watch glass. Exactly four early-stage embryos were mouth-pipetted into 20µL worm lysis buffer per tube using a hand pulled glass capillary. Embryos were then lysed in a thermal cycler (as above) and stored at -80°C.

### Droplet digital PCR

Prior to ddPCR, various sample types were diluted to different degrees in nuclease-free water: sorted-PGC lysates (4X), dissociated-embryo lysates (3000-6000X), whole-embryo lysates (10X), and whole-adult lysates (30 adults lysed in 60µL lysis buffer, 1000X). ddPCR was run according to the manufacturer’s recommendations. Briefly, ddPCR reactions were assembled as 24µL mixes containing 0.1µM of each primer, Bio-Rad QX200 ddPCR EvaGreen Supermix (BioRad 186-4034), 0.1U/µL *Sac*I-HF (New England Biolabs), and 4.8µL of sample. Reactions were incubated in the dark at room temperature for 30-60 minutes to allow *Sac*I-HF (NEB R3156) digestion to linearize/digest DNA prior to droplet generation. After incubation, samples were loaded for droplet generation in a BioRad QX200 Automated Droplet Generator. PCR amplification was performed as follows: 10 minutes at 95°C, 40 cycles of 94°C for 30s and 60°C for 1 minute, followed by 10 minutes at 98°C for all primer pairs. Samples were all run in triplicate, and were immediately analyzed using a BioRad QX200 Droplet reader. All ddPCR reactions were single oligo-pair mixes; therefore, absolute DNA concentrations were calculated using 1D-amplitude plots in BioRad QuantaSoft software.

#### mtDNA copy number quantification by ddPCR

Absolute mtDNA copy number per cell was determined using primer pairs targeting mtDNA (*nd-1)* and gDNA (*cox-4*).

mtDNA –

*nd-1_Fw: 5’- agcgtcatttattgggaagaagac -3’*

*nd1_Rv: 5’- aagcttgtgctaatcccataaatgt -3’*

gDNA –

*cox-4_Fw: 5’- gccgactggaagaacttgtc -3’*

*cox-4_Rv: 5’- gcggagatcaccttccagta -3’*

Two independent ddPCR reactions of the same sample were run simultaneously to determine the mtDNA copies/µL and gDNA copies/µL. mtDNA copy number/cell was calculated as follows: total mtDNAs detected **/** [total gDNA detected **/** (N)], where the ploidy (N)=4 since *C. elegans* PGCs are arrested in the G2 phase of the cell cycle [19]. For L1 feeding experiments the ploidy was calculated based on the expected verses the actual number of gDNAs detected (Fig. S5). Since the ploidy of starved L1 PGCs is constant, we could normalize our data as such. For example, we found that when we sorted 5000 starved L1 PGCs we detected 61 gDNA copies via our ddPCR assay. Therefore, when we sorted 5000 mid-L1 or L2 PGCs and only detected 46 gDNAs we estimated the ploidy as follows:

[(actual copies detected: 46) **/** (expected copies detected: 61)] X 4, where the multiplication factor 4 adjusts the ratio with respect to N=4 for starved L1 PGCs. Thus, for fed L1/L2 PGCs the ploidy (N) can be estimated as approximately 3. This value agrees well with estimated ploidy values based on the calculated cell cycle occupancy times of mitotic germ cells in *C. elegans* adults [57].

#### ΔmtDNA heteroplasmy quantification

mtDNA heteroplasmy was determined using four oligo pairs that specifically detect

*uaDf5, mptDf2,* and their respective complementing WT mtDNAs: For *uaDf5* heteroplasmy –

*uaDf5-mtDNA_Fw: 5’- ccatccgtgctagaagacaaag -3’*

*uaDf5-mtDNA_Rv: 5’- ctacagtgcattgacctagtcatc -3’*

*WT-mtDNA_Fw: 5’- gtccttgtggaatggttgaatttac -3’*

*WT-mtDNA_Rv: 5’- gtacttaatcacgctacagcagc -3’*

For *mptDf2* heteroplasmy –

*mptDf2-mtDNA_Fw: 5’- ggattggcagtttgattagagag -3’*

*mptDf2-mtDNA_Rv: 5’- aagtaacaaacactaaaactcccaac -3’*

*WT-mtDNA_Fw: 5’- cgtgcttatttttcggctgc -3’*

*WT-mtDNA_Rv: 5’- ctttaacacctgttggcactg -3’*

Two independent ddPCR reactions were run simultaneously for each sample to determine the WT mtDNA copies/µL and mutant mtDNA copies/µL. Percent heteroplasmy was then calculated as follows:

[ΔmtDNA **/** (ΔmtDNA + WT mtDNA)] X 100.

### Microscopy

Embryos and larvae were mounted on 5% and 10% agarose pads, respectively. Larvae were immobilized prior to and during image acquisition using 1.25mM levamisole in M9 buffer. Animals were imaged on a Leica SP8 laser-scanning confocal microscope, using a 63X 1.4 NA oil-immersion objective with 488 and 594 nm lasers and HyD detectors; or on a Zeiss AxioImager A2, using a 40X 1.3 NA oil-immersion objective and a CCD camera (model C10600-10B-H, S. 160522; Hamamatsu). For sorted PGC imaging, 5µL of sorted embryonic and larval PGCs in conditioned L-15 (see ‘FACS and PGC isolation’ above) were mounted on custom depression slides to avoid crushing the cells. Sorted PGCs were then imaged on a Zeiss AxioImager A2 as above. Images were analyzed and processed in ImageJ (NIH), and Adobe Photoshop.

### Image analysis

#### Mitochondrial acidification

Acidification of mitochondria was measured in embryos and L1 larvae by determining the ratio of green-to-red fluorescence of mito-mCherry^PGC^ (pKa 4.5) and mito-Dendra2^PGC^ (pKa 6.5). For L1 larvae, 488nm and 594nm laser intensities were adjusted to ensure a similar dynamic range of signal intensity for mito-mCherry^PGC^ and mito-Dendra2^PGC^ within the PGC cell body. Two regions of interests (ROIs) were drawn - one around PGC lobe debris and the other around cell body mitochondria. Red and green signal intensity was then measured and analyzed using ImageJ (NIH) software.

Acidified mitochondria in the embryo were defined as regions of the PGC mitochondrial network where red signal overtook green, such that the measured green-to-red signal ratio was at least two-fold less compared to the greater mitochondrial network (Fig. S7). PGCs of 1.5-fold to 2-fold embryos were imaged and scored categorically as either containing, or not containing, regions of acidified mitochondria. An ROI was then drawn around regions with red dominant signal, and green/red signal intensity was measured in ImageJ. Green/red signal was also measured within an ROI enclosing the rest of the mitochondrial network for comparison.

#### Quantification of mitochondrial localization in PGCs

1.5-fold and 2-fold embryos were imaged as described above. Mitochondrial content was measured as a sum of Mito-Dendra^PGC^ positive voxels within the PGC using ImageJ. A region of interest was then drawn specifically around the PGC cell body using Mem-mCh^PGC^ as a marker, and the fraction of mitochondria in the PGC cell body was calculated as a ratio of total PGC mitochondria.

#### *In vivo* measurement of PGC and whole embryo volume

The volume of PGCs was determined in embryos just prior to lobe formation (bean stage) and in starved L1 larvae. A Z-stack was taken through the PGCs of animals expressing a PGC specific membrane marker (PGC-GFP:: PHPLC1∂1) [58], the volume of both PGCs was measured by defining the PGC surfaces using the image analysis platform Imaris(Oxford Instruments); the volume contained within them was measured and divided by two to determine the volume per single PGC. Three independent biological replicates (N) with sample size n ≥ 18 were used to calculate PGC volume for embryos and L1s. The mean of means was used to calculate the average PGC volume and standard error of the mean (SEM). Embryo volume was calculated by measuring the anterior-posterior and left-right axes of fertilized embryos in ImageJ. Whole embryos were assumed to approximate an ellipsoid, and the volume was calculated using the formula V = 4/3 π a*b*c, where a, b, and c are the radii of the three axes of the ellipsoid (the width and height of embryos were assumed to be equal). Three independent biological replicates (N) with sample size n ≥ 13 were used to calculate whole embryo volume. The mean of means was used to calculate the average PGC volume and standard error of the mean (SEM).

#### Quantification of TFAM foci

Embryos, starved L1, early-L1, mid-L1, late-L1, and L2 larvae were mounted as described above (see ‘Microscopy’). A full Z-stack of the entire germline was taken for each animal. Germline TFAM-GFP/GFP(11) foci were identified using ImageJ to segment TFAM- GFP/GFP11 signal that colocalized with mito-mCherry^PGC^. Colocalized TFAM-GFP/GFP(11) foci were then defined as local signal maxima and counted using the 3D maxima plugin of the ImageJ 3D suite.

#### PGC counts

Embryos and starved L1 larvae were assumed to have exactly two PGCs. For fed larvae expressing TFAM-GFP/GFP(11) and Mito-mCh^PGC^, the number of cells per animal was determined by counting the dark spots in image stacks surrounded by Mito-mCh^PGC^ as a proxy for germ cell nuclei. For cell sorting experiments, fed larvae were mounted and imaged just prior to cell dissociation (see ‘larval cell dissociation’ above), germ cell counts were determined by counting the number of nuclei surrounded by GLH-1-GFP.

#### *Ex vivo* measurement of sorted PGC volume

Sorted embryonic and L1 PGCs were imaged as described above (see ‘Microscopy’).

PGC diameter was calculated by drawing a line across the center of the cell and measuring its length in ImageJ. PGC volume was determined under the assumption that the PGCs approximate a sphere, and volume was calculated with the formula V = 4/3πr^3^.

### Transgene construction

Transgenes *Pmex-5::tomm-20(a.a.1-54)::Dendra2::nos-2^3’UTR^* (plasmid pYA57) and *Pmex-5::gfp1-10::nos-2^3’UTR^* (plasmid pAS07) were constructed by Gibson assembly [59]. Briefly, overlapping primers were used to amplify *tomm-20(a.a.1-54)::Dendra2* to replace *mCherry::moma-1* in *pYA11(Pmex-5:: mCherry::moma-1::nos-2^3’UTR^)*, a derivative of *pCFJ150*. Split *gfp1-10* (based on mammalian versions of the protein) was *C. elegans* codon optimized, designed with introns and ordered as a gBlock (IDT) with overhangs to replace *mCherry-PH* in *pAS06(Pmex-5:: mCherry-PH::nos-2^3’UTR^),* a derivative of *pCFJ150* that lacks a portion the universal MosSCI homology sequence to facilitate CRISPR mediated insertion of the plasmid [60].

### Transgenesis and genome editing

#### MosSCI

*Pmex-5::tomm-20(a.a.1-54)::Dendra2::nos-2^3’UTR^* was microinjected into strain EG8708 to create a single-copy chromosomal insertion on chromosome I via the universal MosSCI method [61].

#### CRISPR/Cas9

In all cases, CRISPR/Cas9 mediated genome editing was performed using pre- incubated Cas9 (Berkeley)::sgRNA (IDT) ribonuclear protein, and injection quality was screened using the co-CRISPR *dpy-10(cn64)* mutation as previously described [62]. DNA repair templates contained ∼25-35bps of homology, and varied depending on the size of insertion as either dsDNA PCR product or plasmid (>150bps), or ssDNA oligos (<150bps) (IDT). crRNAs and insertion sequences are listed in Supplemental Table 2 and Supplemental Sequences. For the generation of putative null alleles, we used the ‘STOP-IN’ method [63] to insert an early stop and frame-shift into either the first, or second, exon of the target gene. All strains generated by CRISPR are included in Supplemental Table 1.

### Statistical analysis and reproducibility

Statistical analysis was performed in GraphPad Prism 9 software. For categorical data, such as scoring acidified mitochondria in PGCs, contingency tables were made and Fisher’s exact test was used to calculate *p*-values. For all other data, two-tailed Student’s *t*-tests (paired and unpaired) were performed. Data in graphs are shown as Superplots [64], with individual data points from three independent color-coded biological replicates (except for ddPCR experiments where small dots are technical replicates of the ddPCR analysis) shown as small dots, the mean from each experiment shown as a larger circle, the mean of means as a horizonal line, and the S.E.M as error bars. Sample size, *t*-test type and *p*-value ranges are reported in Figure Legends. Where applicable, no corrections for multiple comparisons were made to avoid type II errors [65]. For live imaging, embryos and larvae were selected based on orientation on the slide and on health. For all datasets, at least three biologically independent experiments were performed and the arithmetic means of biological replicates were used for statistical analysis.

---

**Figure S1.**
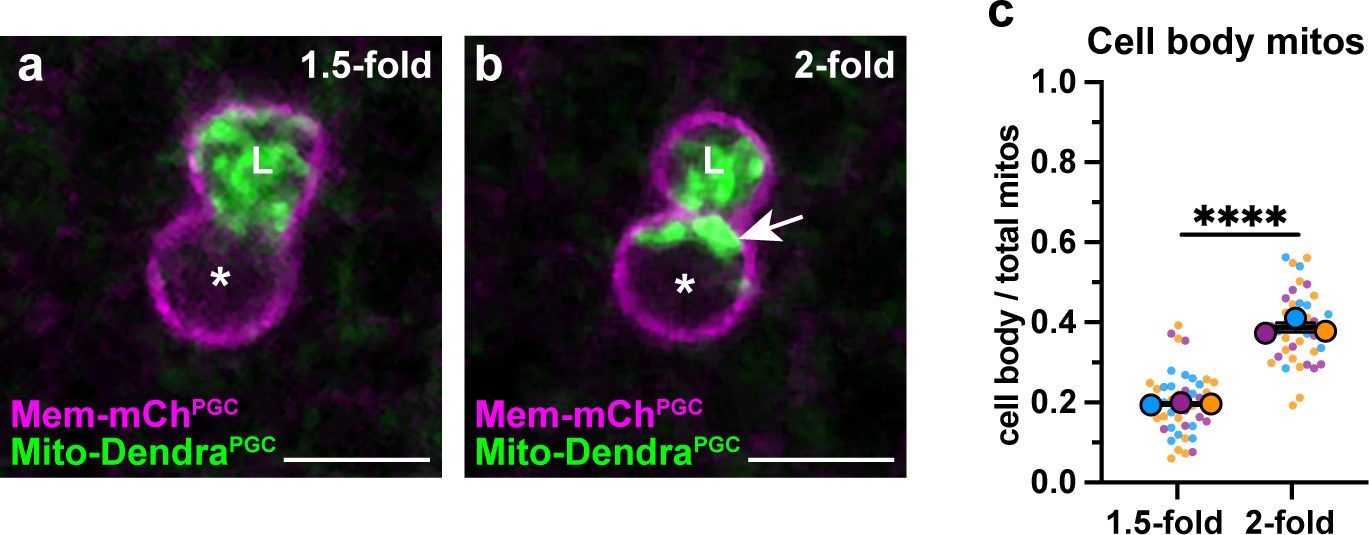
A subset of PGC mitochondria is retained in the cell body prior to lobe digestion **(a-b)** Representative images of plasma membranes and mitochondria in an embryonic PGC as mitochondria localize into lobes (1.5 fold, a) and as lobe cannibalism is initiated (2-fold, b). A subset of mitochondria (arrowhead, b) is retained in the cell body. *, nucleus, ‘L’, lobe. (**c**) Quantification of mitochondrial fraction within the cell body in 1.5-fold and 2-fold stage PGCs. Data in graph: Individual data points from three independent color-coded biological replicates shown as small dots, the mean from each experiment shown as a larger circle, the mean of means as a horizonal line, and the S.E.M as error bars. \*\*\*\**p* ≤ 0.0001, unpaired two-tailed Student’s *t*-test Student’s t-test (g). Scale bars, 5µm.

**Figure S2.**
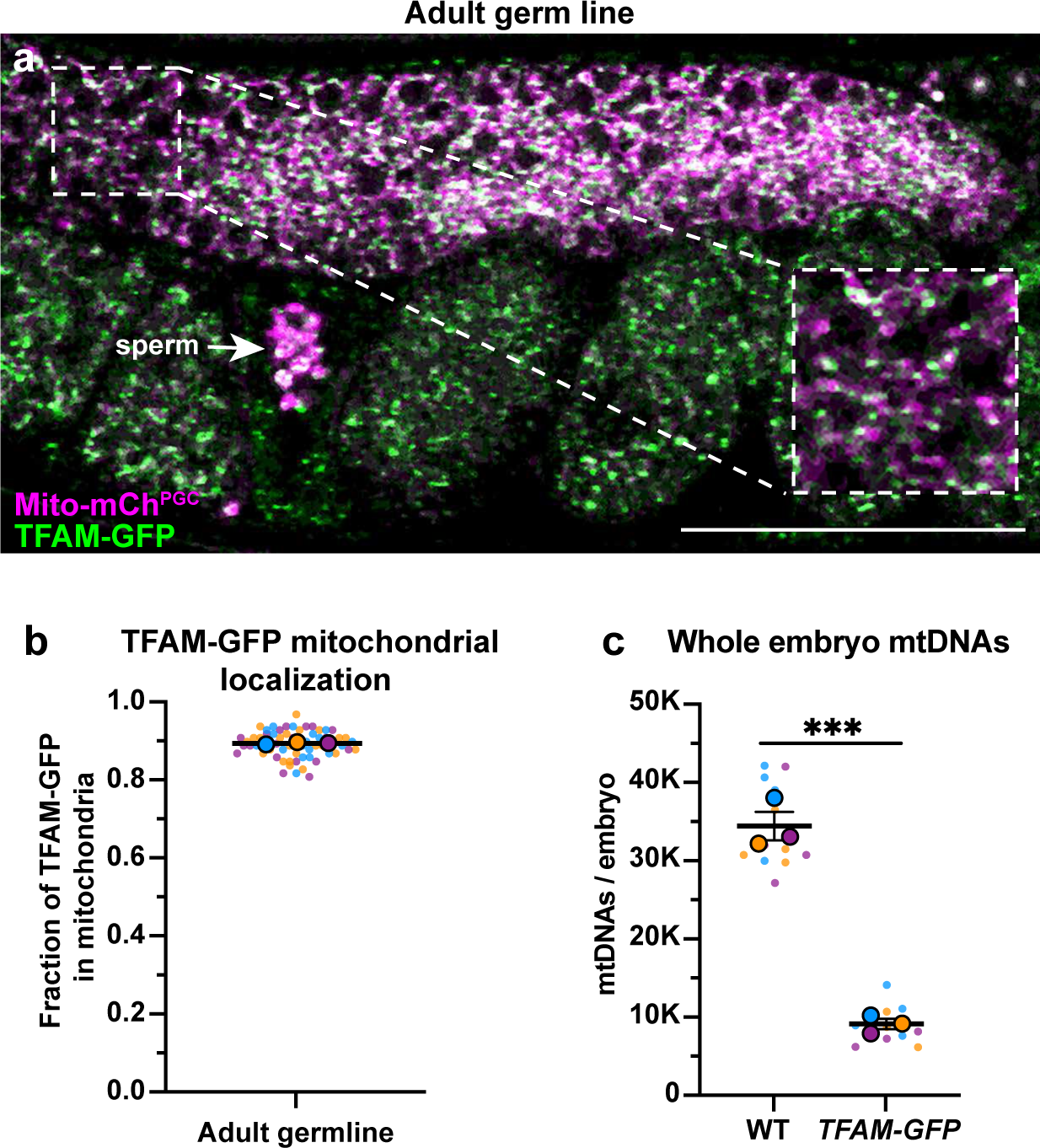
TFAM-GFP mitochondrial localization, and effect on mtDNA copy number (**a**) Endogenously tagged TFAM-GFP and mitochondria in the adult germ line; mitochondria and TFAM-GFP also localize to sperm (arrow). (**b**) Quantification of the fraction of TFAM-GFP overlap with Mito-mCh^PGC^. **(c)** Quantification of mtDNA copy number in wild type and *TFAM-GFP* whole early embryos. Data shown: Small dots are data points from individual worms (b) or technical replicates of ddPCR quantification (c) from each of 3-4 color-coded biological replicates; the mean from each biological replicate is shown as a larger circle, the mean of means as a horizonal line, and the S.E.M as error bars. \*\*\**p* ≤ 0.001, unpaired two-tailed Student’s *t*-test (c). Scale bar, 50µm.

**Figure S3.**
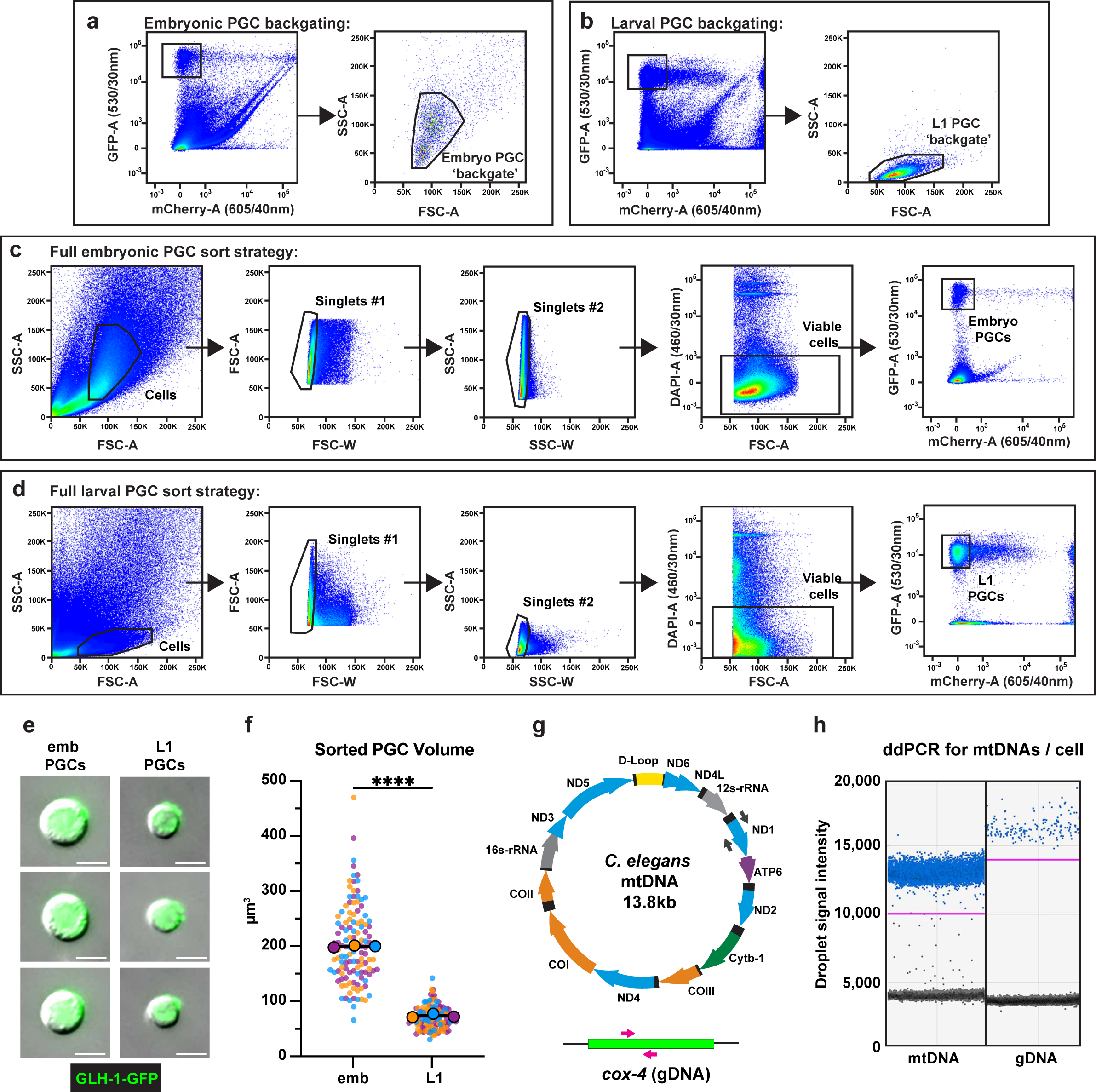
PGC FACS purification gating strategy, quality control, and ddPCR (**a-d**) Full gating strategies for isolating embryonic and L1 PGCs from dissociations. Size exclusion gates (FSC-A x SSC-A) containing PGCs were determined by back-gating on all GFP^+^mCherry^−^ events in embryonic (a) and L1 (b) dissociations. (**c-d**) Following size exclusion, two doublet discrimination gates (FSC-A x FSC-W and SSC-A x SSC-W) were applied to select for singlet cells, DAPI-negative cells were selected for viability, and pure GFP^+^mCherry^−^embryonic (c) and L1 (d) PGCs were sorted. (**e**) Representative images of embryonic and larval PGCs post-FACS. (**f**) Quantification of sorted PGC volume. Small dots are data points from individual cells from each of three color-coded cell sorting experiments; the mean from each sorting experiment is shown as a larger circle, the mean of means as a horizonal line, and the S.E.M as error bars. \*\*\*\**p* ≤ 0.0001, unpaired two-tailed Student’s *t*-test. (**g**) Schematic of *C. elegans* mtDNA and genomic DNA targets, as well as color-coded primer pairs for detecting mtDNA and gDNA by ddPCR. Grey primers, mtDNA, magenta primers, gDNA. (**h**) Representative ddPCR plot for quantifying mtDNA (*nd-1* gene) and gDNA (*cox-4* gene) copy number in sorted PGC lysates. Positive droplets (blue dots), negative droplets (black dots) and threshold for positive droplets (magenta line) are shown. Scale bars, 5µm.

**Figure S4.**
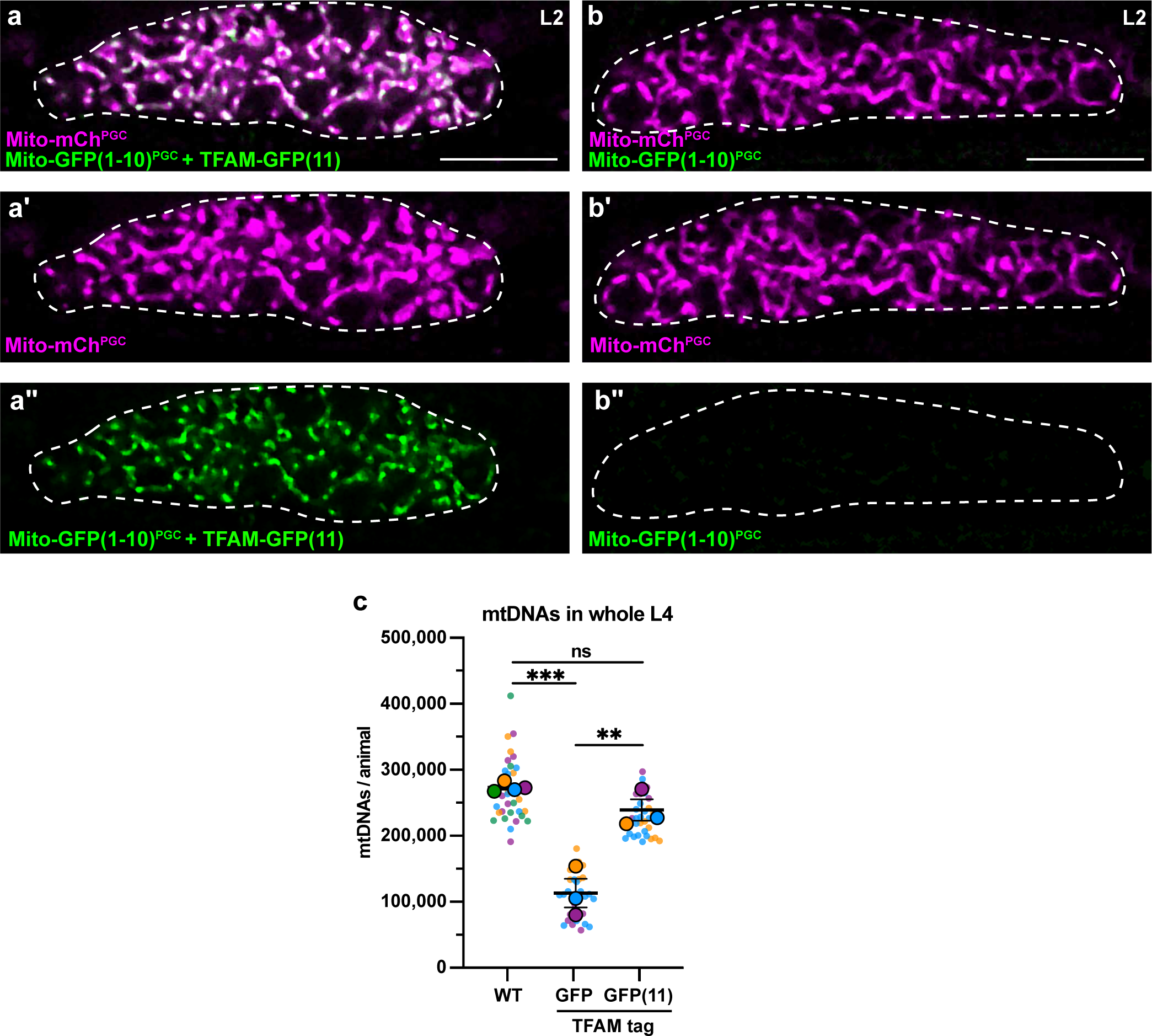
TFAM-GFP(11) visualization and effect on mtDNA copy number (**a-b’’**) Germline mitochondria and GFP(1-10) in L2 larvae, with (a-a’’) or without (b-b’’) endogenously tagged TFAM-GFP(11). Dashed line, outline of gonad. (**c**) Quantification of mtDNA copy number in whole L4 larvae assayed by qPCR in WT, *TFAM-GFP*, and *TFAM-GFP(11); Mito-GFP(1-10)^PGC^* genetic backgrounds. Small dots are data points from individual L4 worms from each of three color-coded biological replicates; the mean from each replicate is shown as a larger circle, the mean of means as a horizonal line, and the S.E.M as error bars. n.s., not significant (*p*> 0.05), \*\**p* ≤ 0.01, \*\*\**p* ≤ 0.001, unpaired two-tailed Student’s *t*-test. Scale bars, 10µm.

**Figure S5.**
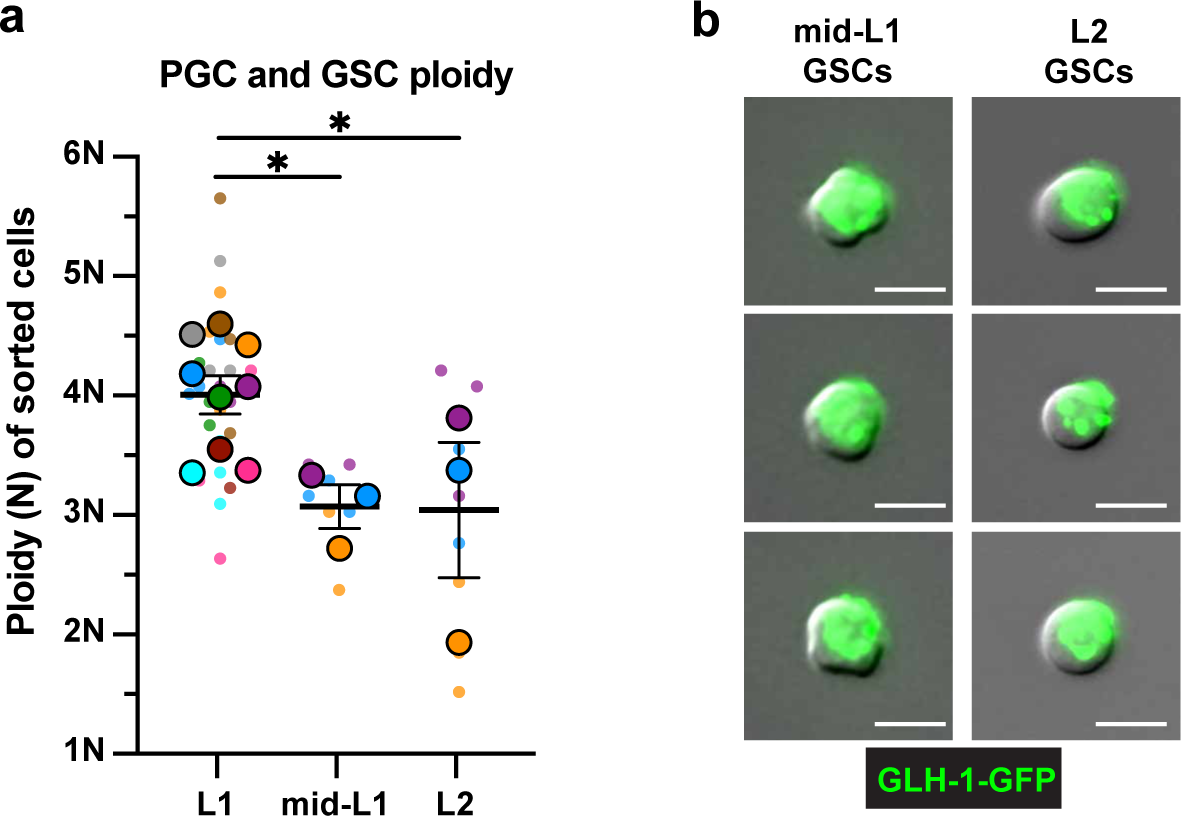
Ploidy and purity of sorted larval GSCs (a) Ploidy of sorted mid-L1 and L2 GSCs (see Methods). Small dots are three technical replicates of ddPCR quantification from each of 3-9 color-coded biological replicates; the technical replicate mean from each experiment is shown as a larger circle, the mean of means as a horizonal line, and the S.E.M as error bars. \**p* ≤ 0.05, unpaired two-tailed Student’s *t*-test. (b) Representative images of mid-L1 and L2 PGCs post-FACS. Scale bars, 5µm.

**Figure S6.**
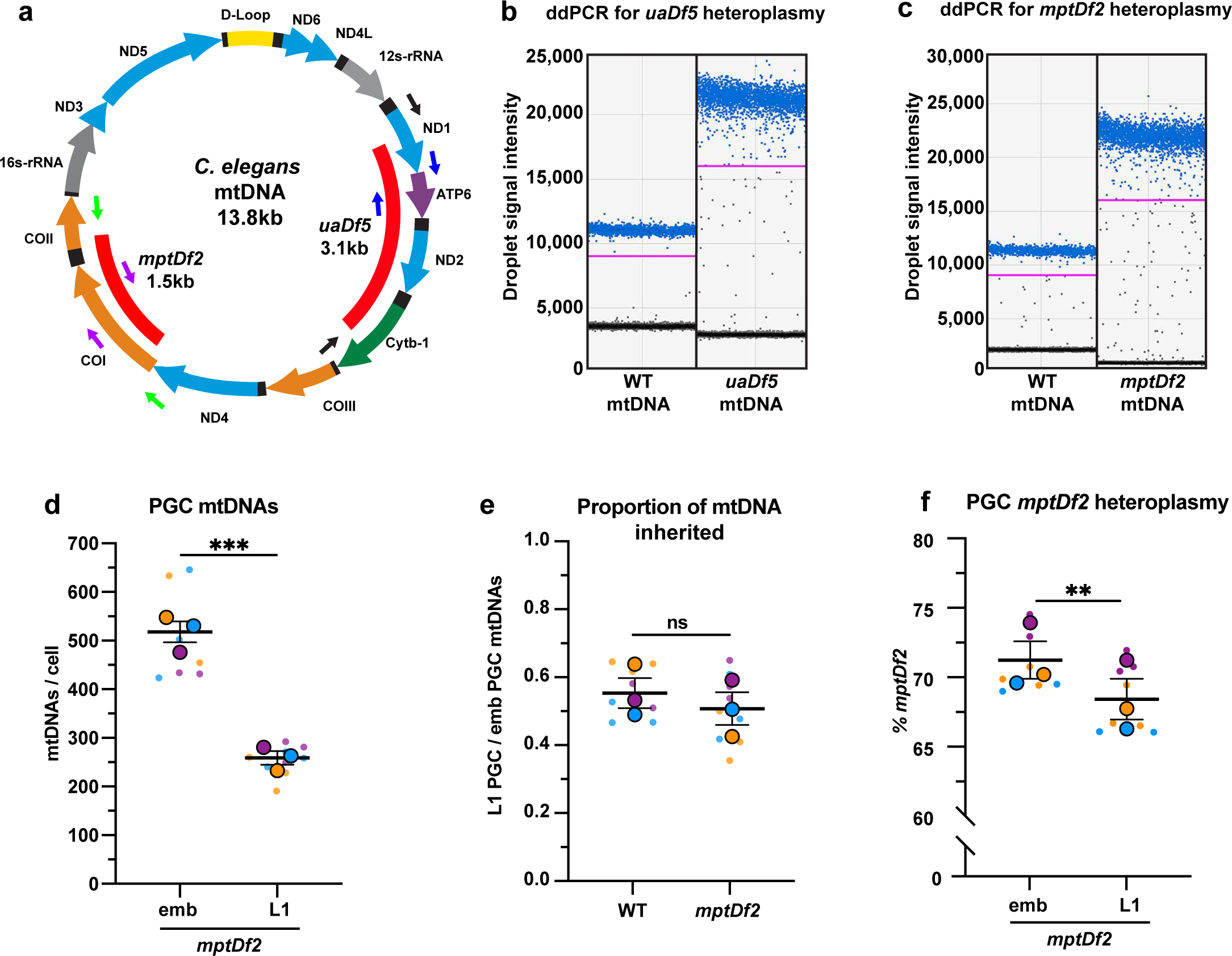
ddPCR primers, detection of mtDNA deletions, and *mptDf2* inheritance in PGCs (a) Schematic of *C. elegans* mtDNA showing mtDNA deletions *uaDf5* and *mptDf2*, as well as color-coded primer pairs for detecting wild-type and mutant mtDNA by ddPCR. Blue primers, wild-type mtDNA (*uaDf5* experiments); black primers, *uaDf5* mtDNA; purple primers, wild-type mtDNA (*mptDf2* experiments); green primers, *mptDf2* mtDNA. (**b**) Representative ddPCR plot for quantifying *uaDf5* and WT mtDNA copy number in sorted PGC lysates. (**c**) Representative ddPCR plot for quantifying *mptDf2* and WT mtDNA copy number in sorted PGC lysates. Positive droplets (blue dots), negative droplets (black dots) and threshold for positive droplets (magenta line) are shown. (**d**) mtDNA copy number in *mptDf2* embryonic and L1 PGCs (**e**) Proportion of *mptDf2* embryonic PGC mtDNAs inherited by L1 PGCs compared to wild type (from data in d and Fig 2b). (**f**) *mptDf2* heteroplasmy in embryonic and L1 PGCs. Data in graphs: small dots are three technical replicates of ddPCR quantification from each of three color-coded biological replicates; the technical replicate mean from each experiment is shown as a larger circle, the mean of means as a horizonal line, and the S.E.M as error bars. n.s., not significant (*p*> 0.05), \*\**p* ≤ 0.01, \*\*\**p* ≤ 0.001 paired (f) and unpaired (e,d) two-tailed Student’s *t*- test.

**Figure S7.**
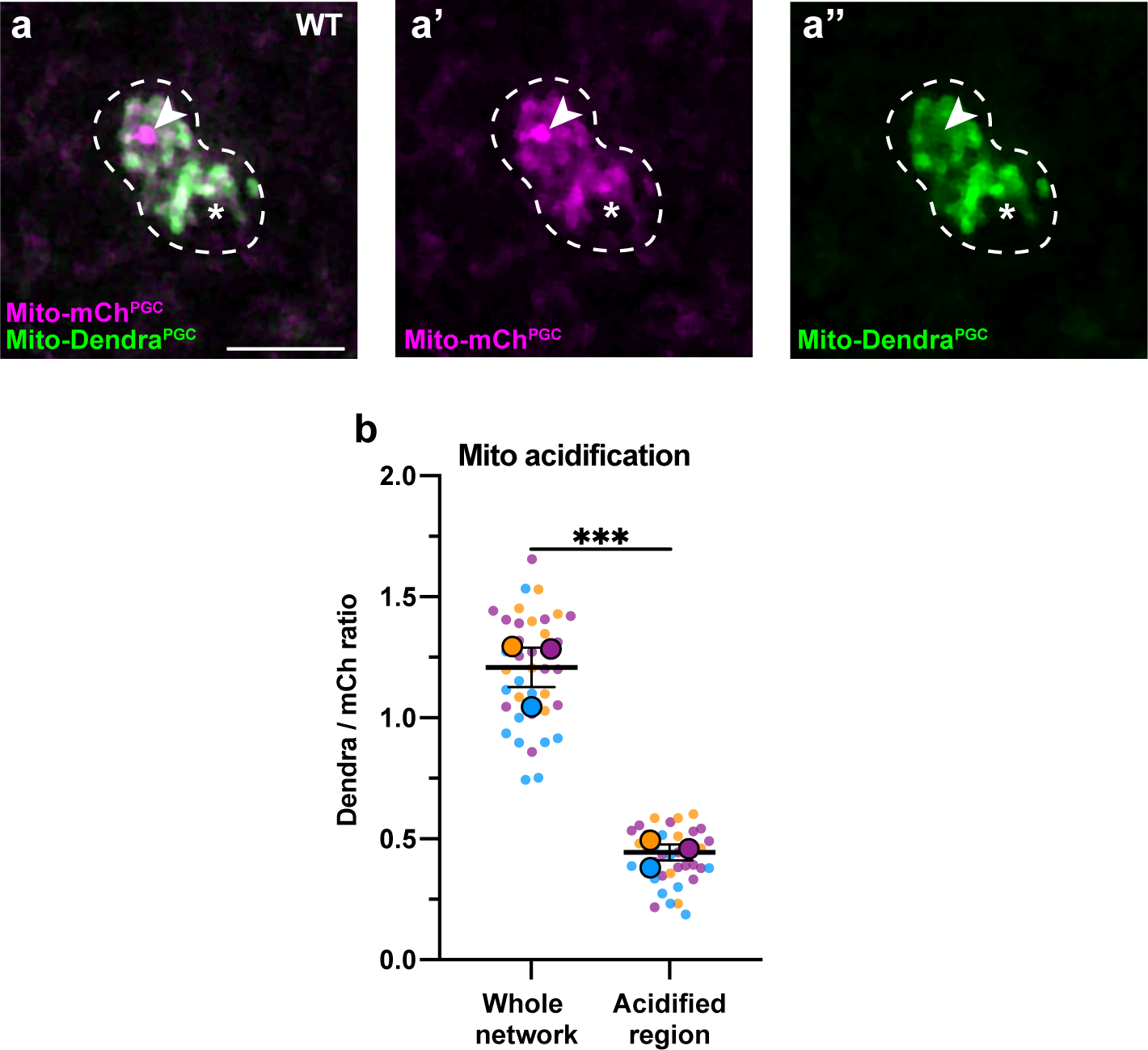
Acidification of a subset of PGC mitochondria (**a-a’’**) Acidified mitochondria (red regions, arrowhead in a, a’) in wild-type PGCs. (**b**) Quantification Dendra / mCherry ratio in whole mitochondrial network and in acidified region. Individual data points from three independent color-coded biological replicates are shown as small dots, the mean from each experiment shown as a larger circle, the mean of means as a horizonal line, and the S.E.M as error bars. \*\*\**p* ≤ 0.001, ratio paired two-tailed Student’s *t*-test. Scale bar, 5µm.

**Supplemental table 1:**
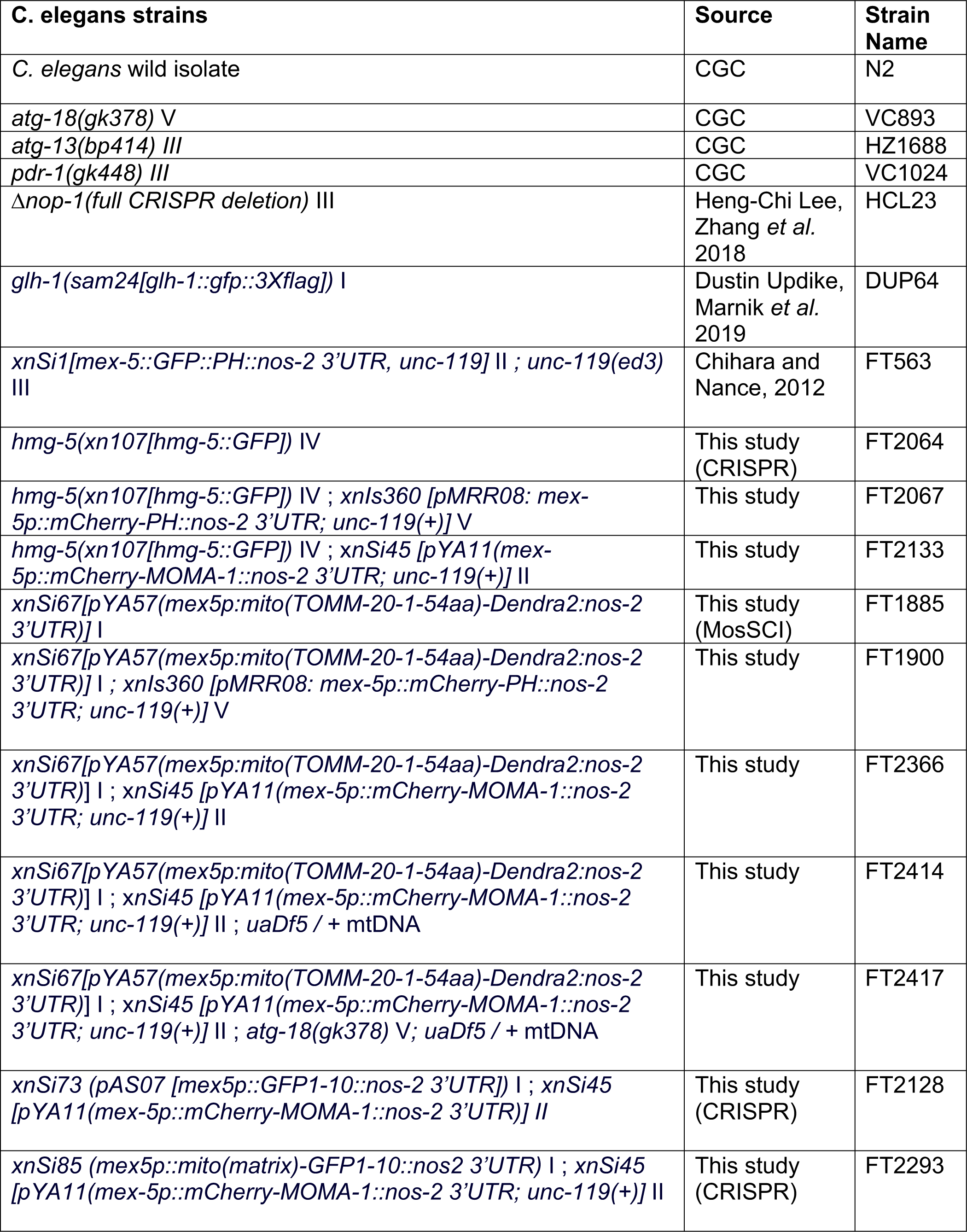
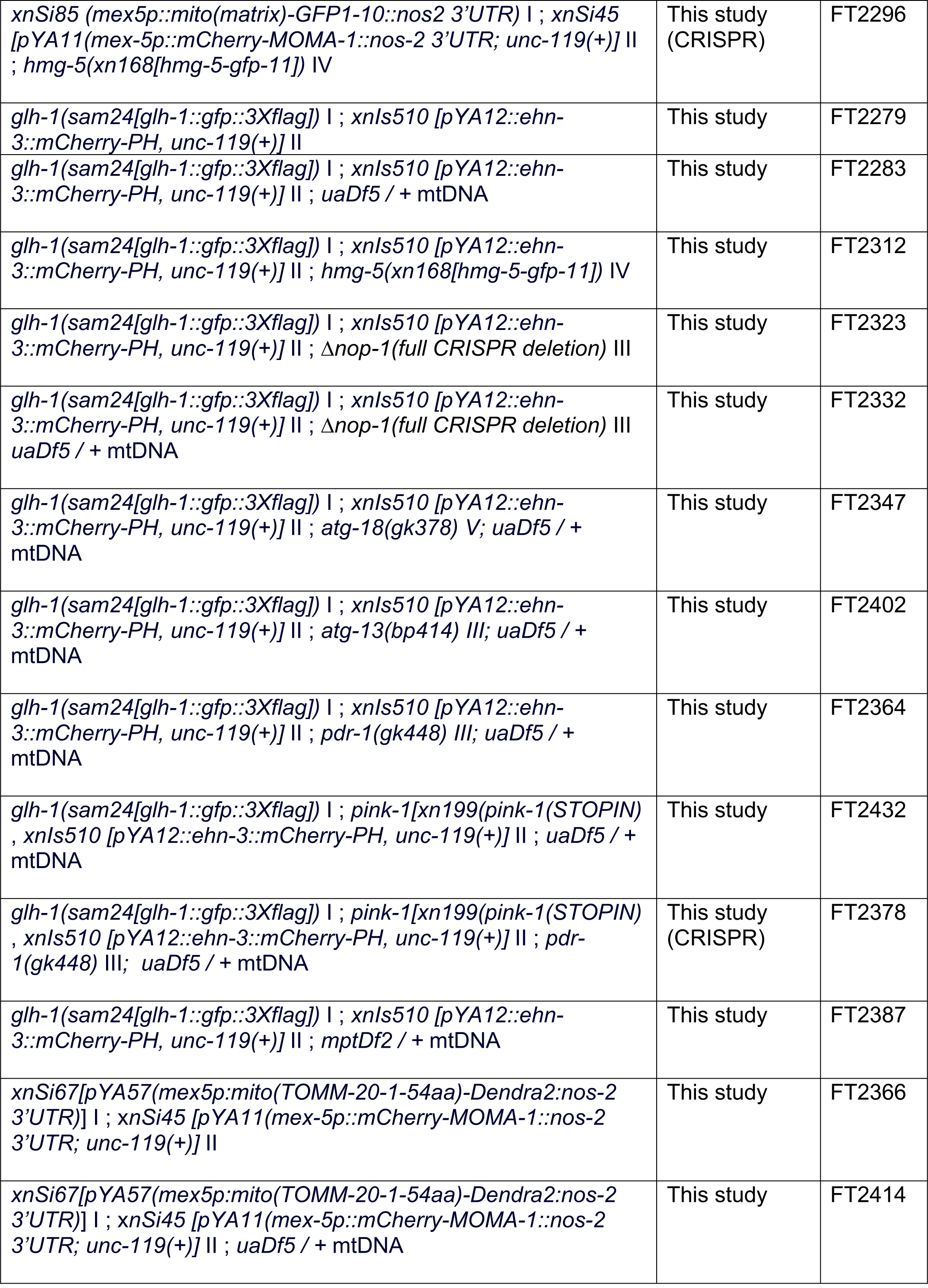
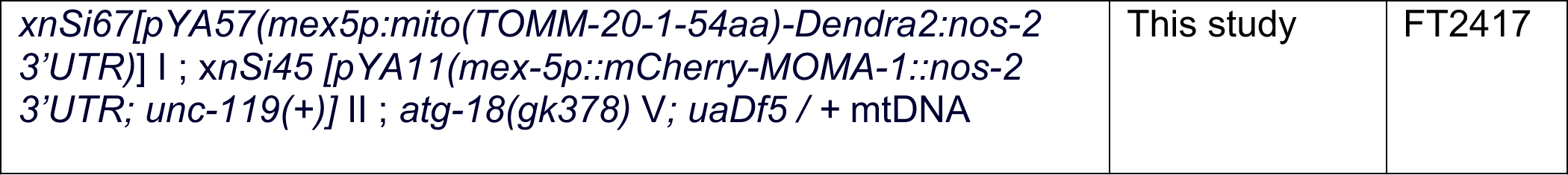

| **C. elegans strains** | **Source** | **Strain Name** |
| --- | --- | --- |
| *C. elegans* wild isolate | CGC | N2 |
| *atg-18(gk378)* V | CGC | VC893 |
| *atg-13(bp414)* III | CGC | HZ1688 |
| *pdr-1(gk448)* III | CGC | VC1024 |
| $\Delta nop-1$ (full CRISPR deletion) III | Heng-Chi Lee, Zhang *et al.* 2018 | HCL23 |
| *glh-1(sam24[glh-1::gfp::3Xflag])* I | Dustin Updike, Marnik *et al.* 2019 | DUP64 |
| *xnSi1[mex-5::GFP::PH::nos-2 3'UTR, unc-119]* II ; *unc-119(ed3)* III | Chihara and Nance, 2012 | FT563 |
| *hmg-5(xn107[hmg-5::GFP])* IV | This study (CRISPR) | FT2064 |
| *hmg-5(xn107[hmg-5::GFP])* IV ; *xnIs360 [pMRR08: mex-5p::mCherry-PH::nos-2 3'UTR; unc-119(+)]* V | This study | FT2067 |
| *hmg-5(xn107[hmg-5::GFP])* IV ; *xnSi45 [pYA11(mex-5p::mCherry-MOMA-1::nos-2 3'UTR; unc-119(+))]* II | This study | FT2133 |
| *xnSi67[pYA57(mex5p:mito(TOMM-20-1-54aa)-Dendra2:nos-2 3'UTR)]* I | This study (MosSCI) | FT1885 |
| *xnSi67[pYA57(mex5p:mito(TOMM-20-1-54aa)-Dendra2:nos-2 3'UTR)]* I ; *xnIs360 [pMRR08: mex-5p::mCherry-PH::nos-2 3'UTR; unc-119(+)]* V | This study | FT1900 |
| *xnSi67[pYA57(mex5p:mito(TOMM-20-1-54aa)-Dendra2:nos-2 3'UTR)]* I ; *xnSi45 [pYA11(mex-5p::mCherry-MOMA-1::nos-2 3'UTR; unc-119(+))]* II | This study | FT2366 |
| *xnSi67[pYA57(mex5p:mito(TOMM-20-1-54aa)-Dendra2:nos-2 3'UTR)]* I ; *xnSi45 [pYA11(mex-5p::mCherry-MOMA-1::nos-2 3'UTR; unc-119(+))]* II ; *uaDf5* / + mtDNA | This study | FT2414 |
| *xnSi67[pYA57(mex5p:mito(TOMM-20-1-54aa)-Dendra2:nos-2 3'UTR)]* I ; *xnSi45 [pYA11(mex-5p::mCherry-MOMA-1::nos-2 3'UTR; unc-119(+))]* II ; *atg-18(gk378)* V ; *uaDf5* / + mtDNA | This study | FT2417 |
| *xnSi73 (pAS07 [mex5p::GFP1-10::nos-2 3'UTR])* I ; *xnSi45 [pYA11(mex-5p::mCherry-MOMA-1::nos-2 3'UTR)]* II | This study (CRISPR) | FT2128 |
| *xnSi85 (mex5p::mito(matrix)-GFP1-10::nos2 3'UTR)* I ; *xnSi45 [pYA11(mex-5p::mCherry-MOMA-1::nos-2 3'UTR; unc-119(+))]* II | This study (CRISPR) | FT2293 |
| *xnSi85 (mex5p::mito(matrix)-GFP1-10::nos2 3'UTR) I ; xnSi45 [pYA11(mex-5p::mCherry-MOMA-1::nos-2 3'UTR; unc-119(+)) II ; hmg-5(xn168[hmg-5-gfp-11]) IV* | This study (CRISPR) | FT2296 |
| *glh-1(sam24[glh-1::gfp::3Xflag]) I ; xnls510 [pYA12::ehn-3::mCherry-PH, unc-119(+)] II* | This study | FT2279 |
| *glh-1(sam24[glh-1::gfp::3Xflag]) I ; xnls510 [pYA12::ehn-3::mCherry-PH, unc-119(+)] II ; uaDf5 / + mtDNA* | This study | FT2283 |
| *glh-1(sam24[glh-1::gfp::3Xflag]) I ; xnls510 [pYA12::ehn-3::mCherry-PH, unc-119(+)] II ; hmg-5(xn168[hmg-5-gfp-11]) IV* | This study | FT2312 |
| *glh-1(sam24[glh-1::gfp::3Xflag]) I ; xnls510 [pYA12::ehn-3::mCherry-PH, unc-119(+)] II ; Δnop-1(full CRISPR deletion) III* | This study | FT2323 |
| *glh-1(sam24[glh-1::gfp::3Xflag]) I ; xnls510 [pYA12::ehn-3::mCherry-PH, unc-119(+)] II ; Δnop-1(full CRISPR deletion) III uaDf5 / + mtDNA* | This study | FT2332 |
| *glh-1(sam24[glh-1::gfp::3Xflag]) I ; xnls510 [pYA12::ehn-3::mCherry-PH, unc-119(+)] II ; atg-18(gk378) V; uaDf5 / + mtDNA* | This study | FT2347 |
| *glh-1(sam24[glh-1::gfp::3Xflag]) I ; xnls510 [pYA12::ehn-3::mCherry-PH, unc-119(+)] II ; atg-13(bp414) III; uaDf5 / + mtDNA* | This study | FT2402 |
| *glh-1(sam24[glh-1::gfp::3Xflag]) I ; xnls510 [pYA12::ehn-3::mCherry-PH, unc-119(+)] II ; pdr-1(gk448) III; uaDf5 / + mtDNA* | This study | FT2364 |
| *glh-1(sam24[glh-1::gfp::3Xflag]) I ; pink-1[xn199(pink-1(STOPIN) , xnls510 [pYA12::ehn-3::mCherry-PH, unc-119(+)] II ; uaDf5 / + mtDNA* | This study | FT2432 |
| *glh-1(sam24[glh-1::gfp::3Xflag]) I ; pink-1[xn199(pink-1(STOPIN) , xnls510 [pYA12::ehn-3::mCherry-PH, unc-119(+)] II ; pdr-1(gk448) III; uaDf5 / + mtDNA* | This study (CRISPR) | FT2378 |
| *glh-1(sam24[glh-1::gfp::3Xflag]) I ; xnls510 [pYA12::ehn-3::mCherry-PH, unc-119(+)] II ; mptDf2 / + mtDNA* | This study | FT2387 |
| *xnSi67[pYA57(mex5p:mito(TOMM-20-1-54aa)-Dendra2:nos-2 3'UTR) I ; xnSi45 [pYA11(mex-5p::mCherry-MOMA-1::nos-2 3'UTR; unc-119(+)) II* | This study | FT2366 |
| *xnSi67[pYA57(mex5p:mito(TOMM-20-1-54aa)-Dendra2:nos-2 3'UTR) I ; xnSi45 [pYA11(mex-5p::mCherry-MOMA-1::nos-2 3'UTR; unc-119(+)) II ; uaDf5 / + mtDNA* | This study | FT2414 |
| *xnSi67*[*pYA57(mex5p::mito(TOMM-20-1-54aa)-Dendra2:nos-2 3'UTR)*] I ; *xnSi45* [*pYA11(mex-5p::mCherry-MOMA-1::nos-2 3'UTR; unc-119(+))*] II ; *atg-18(gk378)* V; *uaDf5* / + mtDNA | This study | FT2417 |

**Supplemental table 2:**
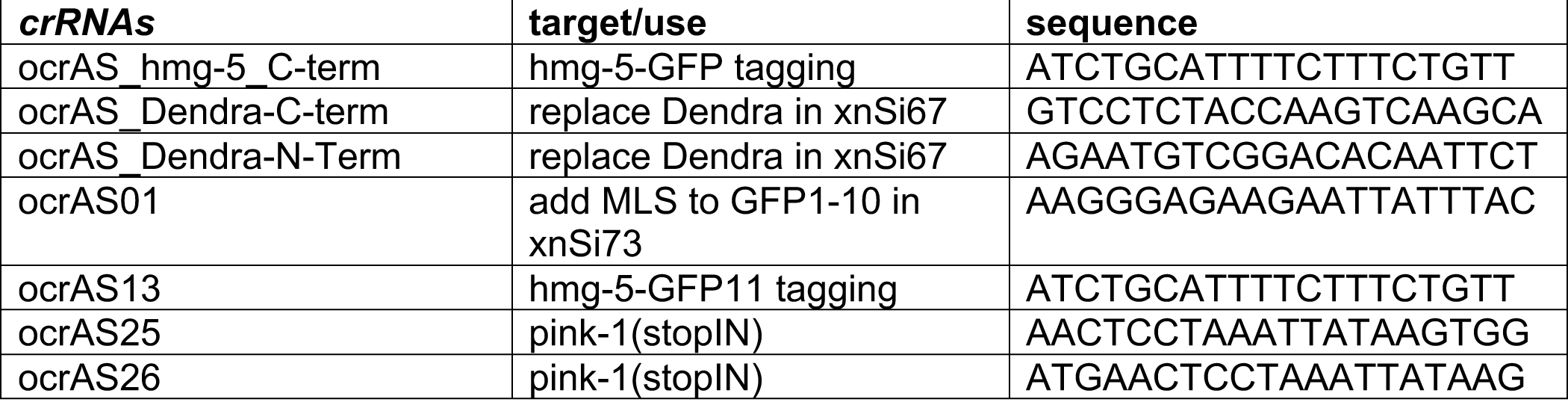

## Supplemental sequences

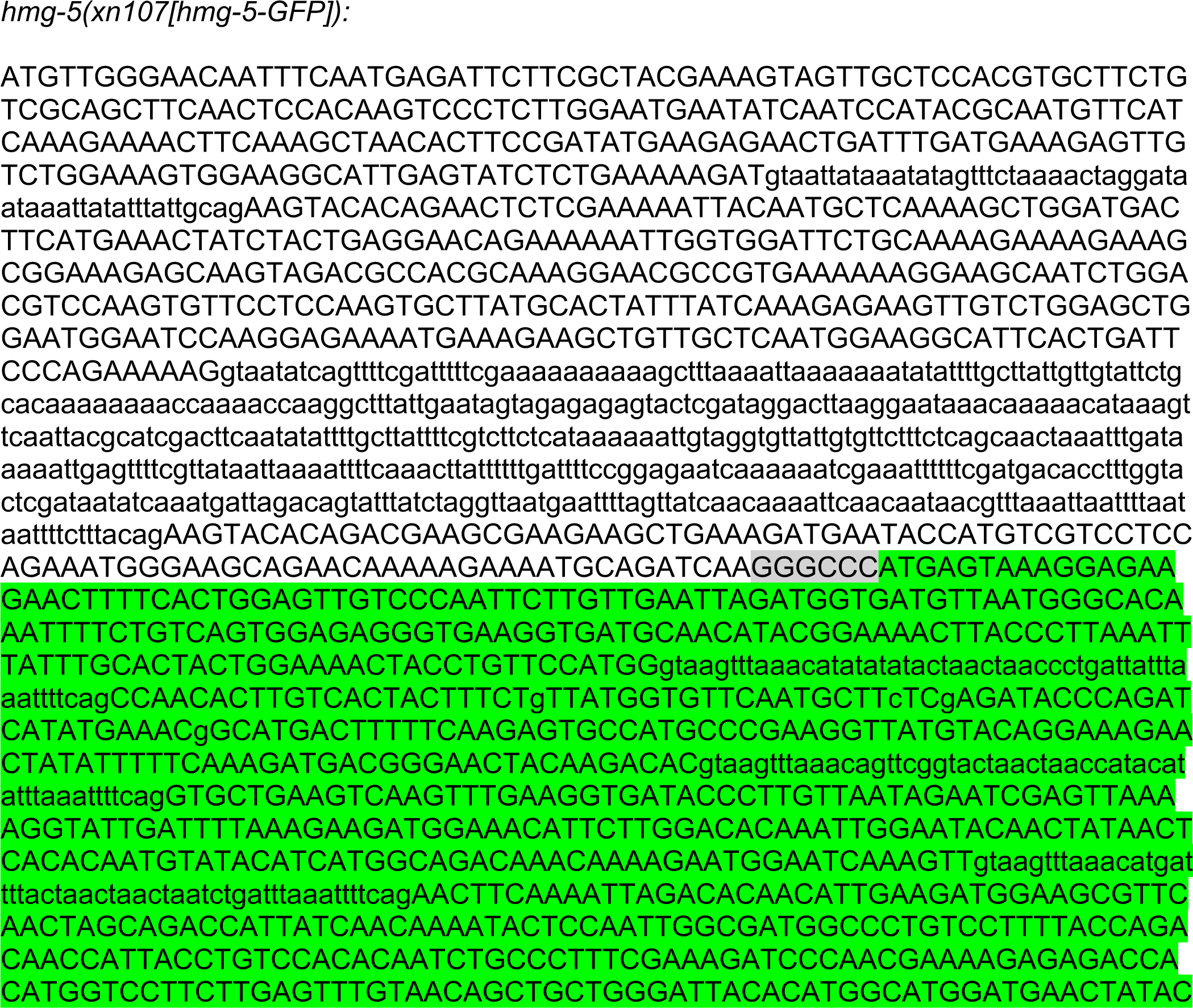
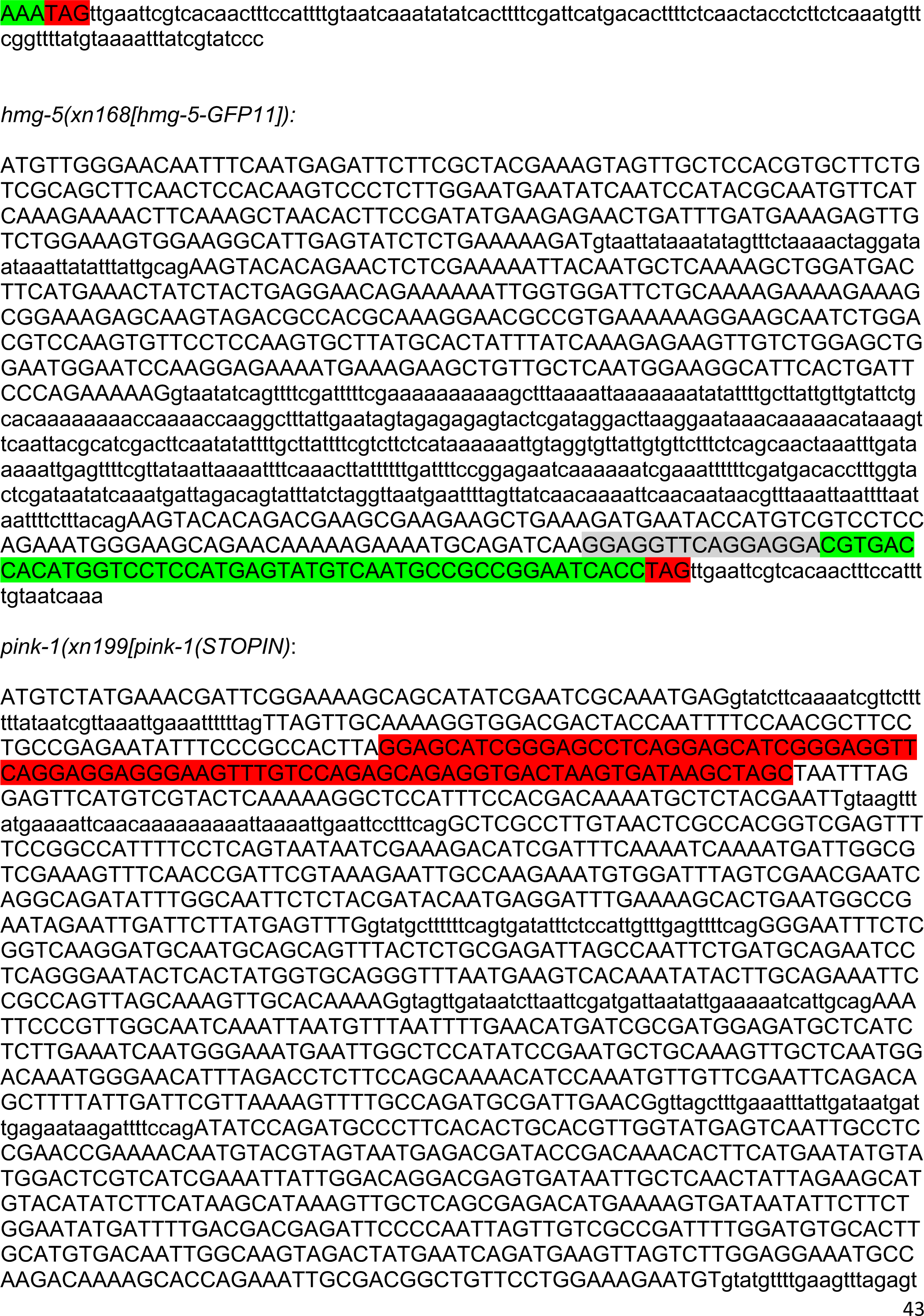
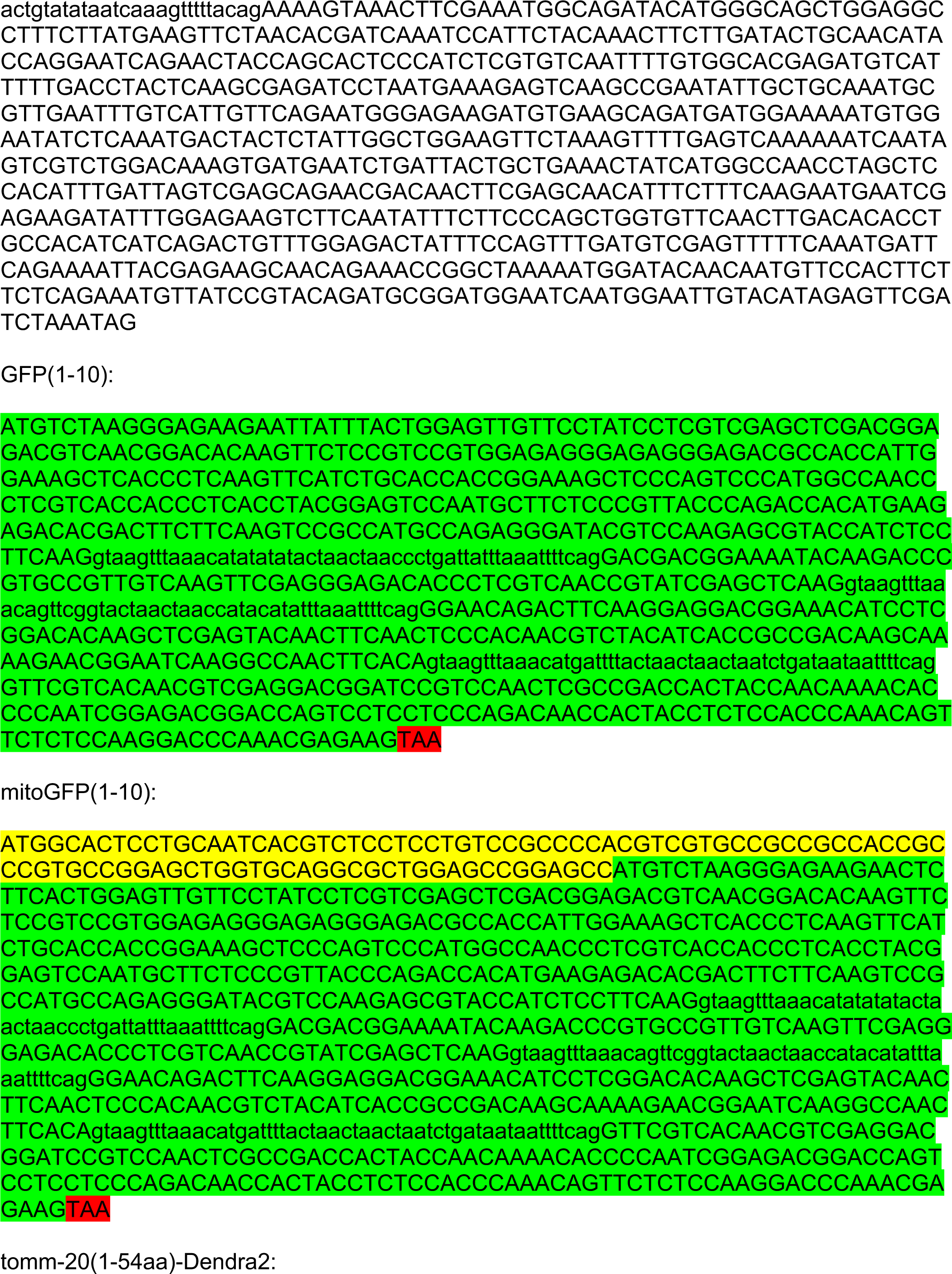
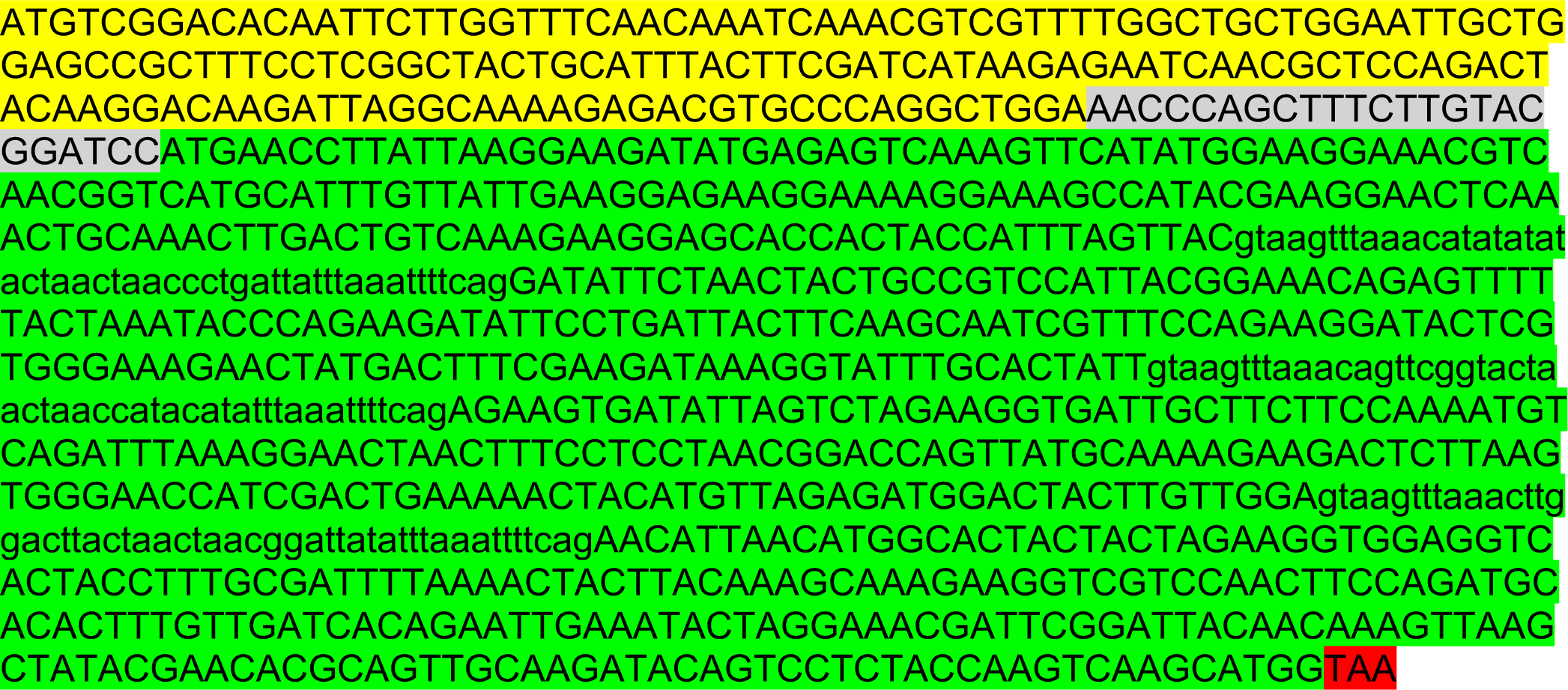

